# Accelerated drug resistant variant discovery with an enhanced, scalable mutagenic base editor platform

**DOI:** 10.1101/2023.10.25.564011

**Authors:** Kristel M. Dorighi, Anqi Zhu, Jean-Philippe Fortin, Jerry Hung-Hao Lo, Jawahar Sudhamsu, Steffen Durinck, Marinella Callow, Scott A. Foster, Benjamin Haley

## Abstract

Personalized cancer therapeutics bring directed treatment options to patients based on the genetic signatures of their tumors. Unfortunately, tumor genomes are remarkably adaptable, and acquired resistance to these drugs through genetic means is an all-too-frequent occurrence. Identifying mutations that promote resistance within drug-treated patient populations can be cost, resource, and time intensive. Accordingly, base editing, enabled by Cas9-deaminase domain fusions, has emerged as a promising approach for rapid, large-scale resistance variant screening in situ. We adapted and optimized a conditional activation-induced cytidine deaminase (AID)-dCas9 system, which demonstrated greater heterogeneity of edits with an expanded footprint compared to the most commonly utilized cytosine base editor, BE4. When combined with a custom sgRNA library, we were able to identify both individual and complex, compound variants in EGFR and BRAF that confer resistance to established EGFR inhibitors. This system and the developed analytical pipeline provide a simple, highly-scalable platform for *cis* or *trans* drug-modifying variant discovery and for uncovering unique insights into protein structure-function relationships.

## INTRODUCTION

Development of resistance to targeted cancer therapies is a common and, unfortunately, anticipated occurrence across a variety of tumor or treatment contexts (1). Among the myriad ways tumors adapt in the presence of a drug, one frequent cause of resistance derives from single nucleotide variants (SNVs), which can reduce drug efficacy through inhibition of target binding, reactivation of signaling pathways, or activation of bypass pathways (2). A well-known example of SNV acquisition is in epidermal growth factor receptor (*EGFR*)-driven non-small cell lung cancer (NSCLC), where the T790M mutation within EGFR, which reduces binding of first generation EGFR inhibitors such as erlotinib or gefitinib, is found in roughly 50% of therapy-insensitive tumors (3, 4). To combat resistance to these molecules and improve overall potency, inhibitors with the ability to co-target the T790M resistance mutation as well as the common activating EGFR variants were developed. However, resistance to these third generation EGFR inhibitors, *e.g.* osimertinib, are being reported in the clinic, for which on-target (*cis*) and non-EGFR (*trans*) SNVs have been observed (5–9). Further, compound mutations that arise independently or layer resistance variants acquired during first line treatment on top of those developed in response to second line therapies are becoming more common (9). The ability to predict therapy-inactivating mutations before drugs under development reach the clinical trial stage could enable the development of treatment options that halt not just one but perhaps several distinct evolutionary paths for a given tumor type. Beyond this, a detailed accounting of the spectrum of mutations that permit drug resistance, whether observed in patients or not, is expected to provide novel insights into intra- or inter-protein functional relationships.

Both gain-of-function (GOF, *e.g.* cDNA expression) and loss-of-function (LOF, *e.g.* CRISPR/Cas9-based pooled knock-out) screening approaches have proven to be effective for identifying drivers of tumor growth/viability and plausible drug resistance mechanisms (10–15). Modification-of-function (MOF) technologies, which rely on various adaptations of CRISPR/Cas9, have emerged as powerful additions to the genetic screening toolbox, by enabling in situ mutagenesis of one or multiple loci at-scale (16, 17). For example, saturation SNV screening of a single *BRCA1* exon was demonstrated with targeted Cas9-induced double-stranded DNA (dsDNA) breaks and subsequent homology-directed repair (HDR) using a battery of oligonucleotide donors (18). More recently, in situ SNV screening has been extended to full-length genes (*NPC1 and BRCA2*) through the application of Cas9-Prime Editing (Cas9-PE), which integrates donor sequences at selected genomic positions by way of custom prime editing guide RNAs (pegRNAs) (19, 20). Despite the robustness of these methods for variant prioritization, HDR or PE-based screening requires pre-designed donor/template sequences. In addition, maximizing the editing efficiency of these methods may require single copy (haploid) target loci or perturbation of specific DNA repair genes within the assayed cell lines. As a result, the throughput and potential for discovering novel variants including compound mutants is limited.

Large-scale variant screens have also been made possible through the direct application of Cas9-Base Editors (Cas9-BEs), which modify DNA sequences through highly efficient chemical reactions (21, 22). This involves the tethering of a deaminase domain selective for cytidines (cytidine base editors, CBEs) or adenines (adenine base editors, ABEs) to a nickase or catalytically-dead Cas9 (Cas9n or dCas9) along with recruitment to a locus of interest using a standard single guide RNA (sgRNA). Once bound to the target site, cytosines or adenines in the editing window (*e.g.* edited bases relative to a given ∼20mer sgRNA target site) will be converted to uracil or inosine, resulting in C>T or A>G edits, respectively (23). Considerable effort has been put into honing the precision (*i.e*. restricting the number of bases modified for each sgRNA) and specificity of Cas9-BEs for use as mutation-correcting therapeutics (24). The predictability of CBE outcomes has also been taken advantage of, with great success, for SNV assessment in a variety of mechanistic and drug resistance screening contexts (25–31). While the precision and restricted editing outcomes of these ABEs or CBEs makes them ideal for determining the phenotypic association of select SNVs, they have limited ability to generate the deep mutational diversity preferred for variant discovery applications. However, more recent studies have described combined ABE/CBEs with the ability to perform simultaneous C>T and A>G edits (32–35) comprising a new set of base editors with higher mutagenic potential.

Here, we sought to develop a simple, high-efficiency toolset for the discovery of single nucleotide and compound variants that promote cancer drug resistance. To do so, we built upon the notion that engineered domains from AID (Activation-Induced cytidine Deaminase), a CBE required for somatic hypermutation during B cell maturation, could be fused or recruited to dCas9 in order to induce C>N or G>N edits at single guide RNA (sgRNA)-adjacent loci (31, 36). Through optimization of a unique AID-dCas9 fusion transgene and its expression parameters, we were able to demonstrate superior mutagenic potential (*e.g.* a variety of editing outcomes at multiple C or G bases within desired loci) and an expanded targeting window compared to similar systems. To explore the capacity for this system to generate functional, compound mutations, we applied it within a tiled sgRNA screen focused on production of coding region mutations in *EGFR* or *BRAF* that convey resistance to distinct, clinically-relevant EGFR inhibitors (erlotinib and osimertinib). Not only were known mutations, such as T790M and T790M/L792F, observed in erlotinib and osimertinib-resistant populations, respectively, but we also discovered several novel drug resistant genotypes, including complex, multi-amino acid variants in BRAF. Computational modeling was used to evaluate the structural impact of select hits. This was followed by validation using transgenic assays across cell lines and drug doses, which enabled us to compare the phenotypic strengths of single vs compound alleles within specific protein domains. Together with our analytical pipeline, the conditional AID-dCas9 system we describe offers a simple, scalable, and effective cell-based platform for novel genetic variant discovery.

## MATERIALS AND METHODS

### Cell culture, transfection

Parental PC-9 and HCC827 cell lines were obtained from the Genentech cell bank (gCell) (37, 38) where they were maintained under mycoplasma-free conditions and authenticated by STR profiling and SNP fingerprinting. Cell lines were cultured in RPMI-1640 growth medium + 10% heat inactivated FBS (Sigma, F4135) + 2mM L-Glutamine and 100 U/mL Penicillin-streptomycin (Gibco, 15140-122). Stable cell lines were generated by transfection of piggyBac transposon (PB) and transposase (PBO, Transposagen/Hera BioLabs) plasmids with GeneJuice (Novogene) using a 10:1 PB:PBO ratio (2ug total DNA) and were selected and maintained in media containing 10 ug/mL blasticidin or 1 ug/mL puromycin. A PC-9.AID-dCas9 clonal cell line was isolated by screening single cell clones for CD81 knockdown by flow cytometry as detailed in the Supplemental Methods.

### Lentiviral production/transduction

sgRNA-expressing and lentiviral packaging plasmids (VSVg/Delta8.9) were transiently co-transfected into HEK293T cells with Lipofectamine 2000. After 72 hours, supernatants were harvested and filtered through a 0.45 uM PES syringe filter. Transduction with lentivirally-encoded sgRNAs was performed in 8 ug/mL polybrene with direct addition of filtered supernatants. Transduced cells were selected with 1 ug/mL puromycin to select for sgRNA-expressing cells.

### Vector sequences

Base editor and *EGFR/BRAF* transgene sequences are detailed in the Supplemental Methods. Single guide RNA sequences are included in the Supplemental Data. piggyBac plasmids were obtained from Transposagen/Hera Biolabs, pLKO lentiviral vectors were obtained from Sigma (SHC001) and cloning was performed by GenScript.

### sgRNA tiling library design

To generate an sgRNA library targeting *EGFR*, *BRAF* and *ESR1*, we designed all possible sgRNAs targeting exons of the canonical Ensembl transcripts (ENST00000275493, ENST00000646891 and ENST00000206249, respectively) using the crisprVerse Bioconductor ecosystem (Hoberecht et al., 2022). Ensembl release 103 (genome build GRCh38) was used. Only protospacer sequences adjacent to a canonical SpyCas9 PAM sequence NGG were considered. This resulted in 785 sgRNAs for *EGFR*, 311 sgRNAs for *BRAF* and 367 sgRNAs for *ESR1* (see Supplemental Data for complete sgRNA list). We added a set of 47 non-targeting controls (NTCs) consisting of random sequences not found in the human genome to serve as negative control gRNAs. The sequences were produced in an arrayed oligo synthesis pool (Cellecta, Inc) and sub-cloned into a puromycin selectable lentivirus vector pLKO_SHC201 (Sigma).

### BE sgRNA tiling screen

The lentiviral sgRNA tiling library consisting of 1510 sgRNAs was transduced into clonal AID-dCas9.PC-9 cells (MOI of 0.3) to achieve 1000x coverage of each sgRNA and selected with 1 ug/mL Puromycin 72 hrs later. Two replicate flasks of cells were split every 3 days and cultured with 2ug/mL doxycycline (DOX) for 0, 3, 6 or 9 days, before seeding cells in 40 nM erlotinib, 200 nM erlotinib, 20 nM osimertinib, 100 nM osimertinib or DMSO. DMSO treated cells were collected after 3 passages over the course of 9 days, while drug treated cells were collected as resistant colonies emerged and were expanded. All samples are present in duplicate, with the exception of two samples (100nM osimertinib D0 DOX replicate B and 100nM osimertinib D3 DOX replicate A) which were lost during sample preparation and excluded from the analysis.

### Genomic DNA isolation and PCR amplification

For the pooled tiled sgRNA screen, genomic DNA was isolated with the Gentra Puregene Cell Kit (158788, Qiagen). DNA concentration was measured using the Qubit dsDNA BR Assay (Q32853, Molecular Probes). To quantify sgRNA abundance, sgRNA amplicons were generated by PCR of 2 ug genomic DNA per 100 ul reaction with Q5 2X Hot Start Master Mix (M0494L, NEB) and enough reactions to maintain 1000x screening sgRNA representation. PCR primers targeting all sgRNAs (5’ - tcttgtggaaaggacgaggtaccg - 3’ and 5’ - tctactattctttcccctgcactgt - 3’) were used with the following conditions: Denaturation for 3 min at 98 °C, followed by 28 cycles of thermal cycling at 98 °C for 10 sec, 64.5 °C for 30 sec and 72 °C for 20 sec. To identify drug resistant mutations, genomic DNA isolated from the tiling screen was amplified using the following primers: *EGFR* region 1 (5’ - atgcgtcttcacctggaagg - 3’ and 5’ - gacatagtccaggaggcagc - 3’), *EGFR* region 2 (5’ - tccaggaagcctacgtgatg - 3’ and 5’ - cccgtatctcccttccctga - 3’) and *BRAF* region 1 (5’ - gcgaacagtgaatatttcctttgat - 3’ and 5’ - acgggactcgagtgatgattg - 3’) using PCR conditions identical to those above, except primer annealing was performed at 58°C. For assaying of base editing in an arrayed format, genomic DNA was isolated with the Quick-DNA 96 Kit (Zymo Research, D3011). Genomic regions in *CD81* and *OR5M9* were amplified using the following primers: *CD81* (5’ - ccccctgtgcatgtgacc - 3’ and 5’ - accgtctcgtggaaggtct - 3’) and *OR5M9* (5’ - ggtgtaaaacacagccaccatt - 3’ and 5’ - acatattccctctcggtggtc - 3’) and identical PCR conditions to those described above, except primer annealing was performed at 60°C. All PCR products were purified with AMPure XP Reagent (A63881, Beckman Coulter) using standard protocols.

### Read alignment and statistical analysis for pooled sgRNA amplicon sequencing

Amplicons were sequenced on an Illumina HiSeq2500 instrument, using 50 bp paired-end reads. For each read pair, both read sequences were searched for an exact match of any one of the gRNA barcode sequences, using the screenCounter R package available in the crisprVerse ecosystem (39). The number of reads matched to each guide in each sample was used to obtain a guide-by-sample count matrix for further analysis. Raw count data were stored in a standard Bioconductor Summarized Experiment object (40). Normalization is needed to adjust for the difference in sequencing depth between samples, and to account for potential compositional biases due to the competitive nature of pooled genetic screening. We estimated normalization factors for each sample by applying the TMM method (41). We performed a differential abundance analysis at the gRNA level using the limma-voom approach (42). Specifically, we fitted a linear model to the log-CPM values for each gRNA, using voom-derived observation and quality weights. We performed robust empirical Bayes shrinkage to obtain shrunken variance estimates for each gRNA, and we used moderated F-tests to compute p-values for each of the two-group comparisons of interest. To control the FDR in each comparison, we applied the standard Benjamini-Hochberg method (43) to obtain an adjusted p-value for each gRNA.

### Read alignment and statistical analysis for arrayed bulk amplicon sequencing

Amplicons were sequenced on an Illumina MiSeq using 150 bp or 200 bp single-end reads. Sequencing reads were aligned to the genome (GRCh38) and mutations were called using a previously described CRISPR editing analysis workflow (44). Mutations occurring within a 50 bp window centered on the 5’ end of the sgRNA were counted, alleles with more than one read were included in the analysis and an allele frequency (percent of total reads per genomic region per sample) was computed for each allele identified. Amino acid changes were annotated using the Ensembl Variant Effect Predictor (VEP) (45). Percent editing at each nucleotide was calculated by splitting alleles into individual mutations and summing the associated allele frequencies. Editing percentages were smoothed across nucleotide positions for each mutation separately using a LOWESS curve (smoother span value of 0.3 for C>T, C>A, C>G, G>A, G>C, G>T). Relative editing weights between 0 and 1 were calculated by dividing all editing percentages by the maximal editing percentage value taken across all possible mutations. To identify enriched alleles from bulk amplicon-sequencing of *EGFR* region 1 and 2 and *BRAF* region 1 in drug samples compared to DOX samples, we fit an empirical Bayesian negative binomial regression model for each detected allele in the experiment. To increase the statistical power, samples of different time points were pooled within respective drug concentrations or DOX and DMSO. The raw count of reads in the allele-by-sample matrices were normalized by sample and region specific total reads, and the change of normalized read counts for each allele were modeled against the drug condition (per drug concentration versus DOX). Due to the low sample size and the sparsity of read counts in DOX samples (baseline), we employed the empirical Bayesian framework to stabilize the estimates and reduce overall error in inference. The dispersion parameter in the negative binomial distribution of normalized counts was estimated with the DESeq2 (46) framework where default parametric trend prior for dispersion and mean relationship was used. The Log2 Fold Change (LFC) of differential allele counts in drug versus DOX samples was estimated as the maximum a posteriori from using an empirical Cauchy prior distribution (47), where the scale of the Cauchy prior is estimated from maximum likelihood estimates of the LFCs. The associated s value (48) was calculated and a cut off of 0.01 was used to control the error rates of the wrong sign of estimates (49).

### Validation screen and single cell DNA sequencing

AID-dCas9.PC-9 cells were infected at 0.3 MOI with the pooled sub-library containing 12 sgRNAs identified as top hits in the tiling screen. Following selection with 1 ug/mL Puromycin and treatment with DOX for 6 days, cells were treated with 100 nM erlotinib or 20 nM osimertinib, changing media every 3 days until resistant cell colonies emerged and expanded between 15-18 days later. Cells were frozen, shipped to Mission Bio and samples processed utilizing the Mission Bio Tapestri^®^ Platform to perform targeted high-throughput single cell DNA sequencing using materials, equipment, supplies and methods described in the Mission Bio Tapestri^®^ User Guide (https://support.missionbio.com/hc/en-us/articles/360045396514-Tapestri-Single-cell-DNA-Sequencing-V2-User-Guide). In brief, after resuspending single cells in Mission Bio cell buffer at 3-4, 000 cells/µL, the cells were put into the Mission Bio Tapestri^®^ cartridge, where single cells are individually partitioned into nano-droplets with lysis buffer (Mission Bio) and incubated at 50°C for 60 minutes on the PCR block to lyse the cells and release the DNA from the chromatin. The DNA is then re-loaded into the Tapestri^®^ cartridge, where barcoding beads and PCR reagents are combined in a second merged encapsulation. UV light is applied for 10 minutes to release the barcoded DNA from the beads, and the DNA is amplified via multiplexed PCR within the droplets (Mission Bio). The droplet emulsions are then broken, and the DNA is extracted and purified using Ampure XP Beads (0.72X). Qubit Fluorescence Quantification (Invitrogen) was used to quantify the concentration of DNA at an expected range of 0.2-4 ng/µL. The multiplexed primer panel used in this study was custom designed by Mission Bio to amplify 150-250 bp centered on the targeting sites of sgRNAs included in the sub-library. For library construction, the i5 and i7 indexes (Illumina) are added, and library amplification is performed on the PCR block. The library is then purified using Ampure XP beads (0.69X) for an expected on-target size range of 350-550 bp. Qubit is used as a preliminary raw quantification of DNA concentration, followed by the BioAnalyzer (Agilent) for a more detailed view of the fragment size distribution in addition to the targeted concentration. The libraries were then diluted to 5 nM (0.9-1.3 ng/µL) and sequenced on the Illumina Nextseq550 High Output at 150 bp paired end, following Illumina protocols. Samples were then demultiplexed using the Mission Bio Tapestri Pipeline for computational analysis.

### Mutation enrichment analysis from Tapestri data

Barcoded single cell reads were analyzed using the CRISPR editing analysis workflow described above. After reducing noise, data was processed in two different ways. To determine amino acid changes enriched in drug treatment vs DOX, cells containing amino acid changes were counted, data was normalized by total cells per amplicon per sample and was fitted using an empirical Bayesian negative binomial regression model. To identify enriched alleles in drug treatment vs. DOX, we used sample-level pseud-bulk data, by summing read counts across cells within each sample. These pseudo-bulk allele counts were then fitted using an empirical Bayesian negative binomial regression model after normalization by total reads per amplicon per sample. More detailed methods for single cell read processing and mutation enrichment analysis can be found in the Supplemental Methods.

### CellTiter-Glo

Cells were seeded at a density of 2, 000 cells per well in 96-well plates 24 h prior to treatment. Cells were treated with a dose titration of erlotinib or osimertinib either in the presence of DOX or DMSO (control). After 3 days of treatment, cell viability was measured using CellTiter-Glo Luminescent Cell Viability Assay (Promega, G7570) following the manufacturer’s protocol. Reagents and plates were equilibrated to room temperature.

### Western Blotting

500, 000 cells/well were seeded into a 6-well plate (Corning, 3516). The following day, cells were subjected to the following media conditions: 2 ug/mL DOX or DMSO (as a control) and either 200 nM erlotinib or 20 nM osimertinib. After 48 hr incubation, cells were harvested and lysed in NP-40 lysis buffer (bioWORLD, 22040045-2) for 15 min on ice. Lysates were cleared by centrifugation, protein concentration measured by BCA (Thermo Fisher, 23225) and prepared for SDS-PAGE using NuPAGE LDS Sample Buffer (4×) (Invitrogen, NP0007) and NuPAGE LDS Sample Reducing Agent (10×) (Invitrogen, NP0009). 20 ug of the boiled samples were loaded onto a NuPAGE 4-12% BisTris gel (Invitrogen, WG1402BOX) and afterwards transferred to nitrocellulose using the Trans-Blot Turbo system (BioRad, 1704150). Blots were stained with the following primary antibodies diluted in Intercept (TBS) Blocking Buffer (LiCor, 927-60001): mouse anti-BRAF (Santa Cruz Biotech, sc-5284, 1:200), mouse anti-ERK (Cell Signaling, 9107, 1:1000), rabbit anti-pERK (Cell Signaling, 9101, 1:1000), mouse anti-MEK (BD Biosciences, 610122, 1:1000), rabbit anti-MEK (Cell Signaling, 9121, 1:1000), mouse anti-EGFR (MBL, MI-12-1, 1:1000), rabbit anti-pEGFR (Cell Signaling, 3777, 1:1000), and mouse anti-β-actin (LiCor, 926-42212, 1:1000). Secondary antibodies were IRDye-conjugates from LiCor (IRDye 800CW donkey anti-rabbit IgG, 926-32213, 1:5000); IRDye 680RD donkey anti-mouse IgG, 926-68072, 1:5000) and blots analyzed by ChemiDoc MP fluorescence imaging (Biorad, 12003154). Images were processed using Image Lab software (BioRad).

### Figure production

Plots and graphs were generated using ggplot2 and custom scripts in R, Prism 9 (GraphPad software) or the PyMOL Molecular Graphics System, Version 2.0 Schrödinger, LLC. All figures were assembled and annotated in BioRender.

## RESULTS

In order to enhance the mutagenic capacity of a commonly used CBE, BE4, we incorporated a series of modifications (Fig 1A, B). First, we decided to replace rApobec1 with a directly-fused hyperactive AID (AID*delta) cytidine deaminase. AID has a wider editing window and reduced sequence preference compared to Apobec1 (17, 23) and has been successfully utilized for mutagenic screening, in which recruitment of AID*delta was achieved via an MS2-MCP system, yielding low efficiency and highly dispersed edits (31). By fusing AID*delta to Cas9n, we aimed to increase the efficiency and concentration of edits on the target locus. Additionally, we removed the tandem Uracil DNA Glycosylase Inhibitor (UGI) domain that prevents excision of uracil, in order to promote error prone DNA repair pathways and increase the mutational spectrum (50, 51). Finally, we compared Cas9 nickase (Cas9n), which has been shown to increase editing efficiency in context with BEs (50), to a nuclease-deficient version of Cas9 (dCas9), used previously in C-terminal fusions with AID and PmCDA1, an AID orthologue from sea lamprey (36, 52).

**Figure 1.**
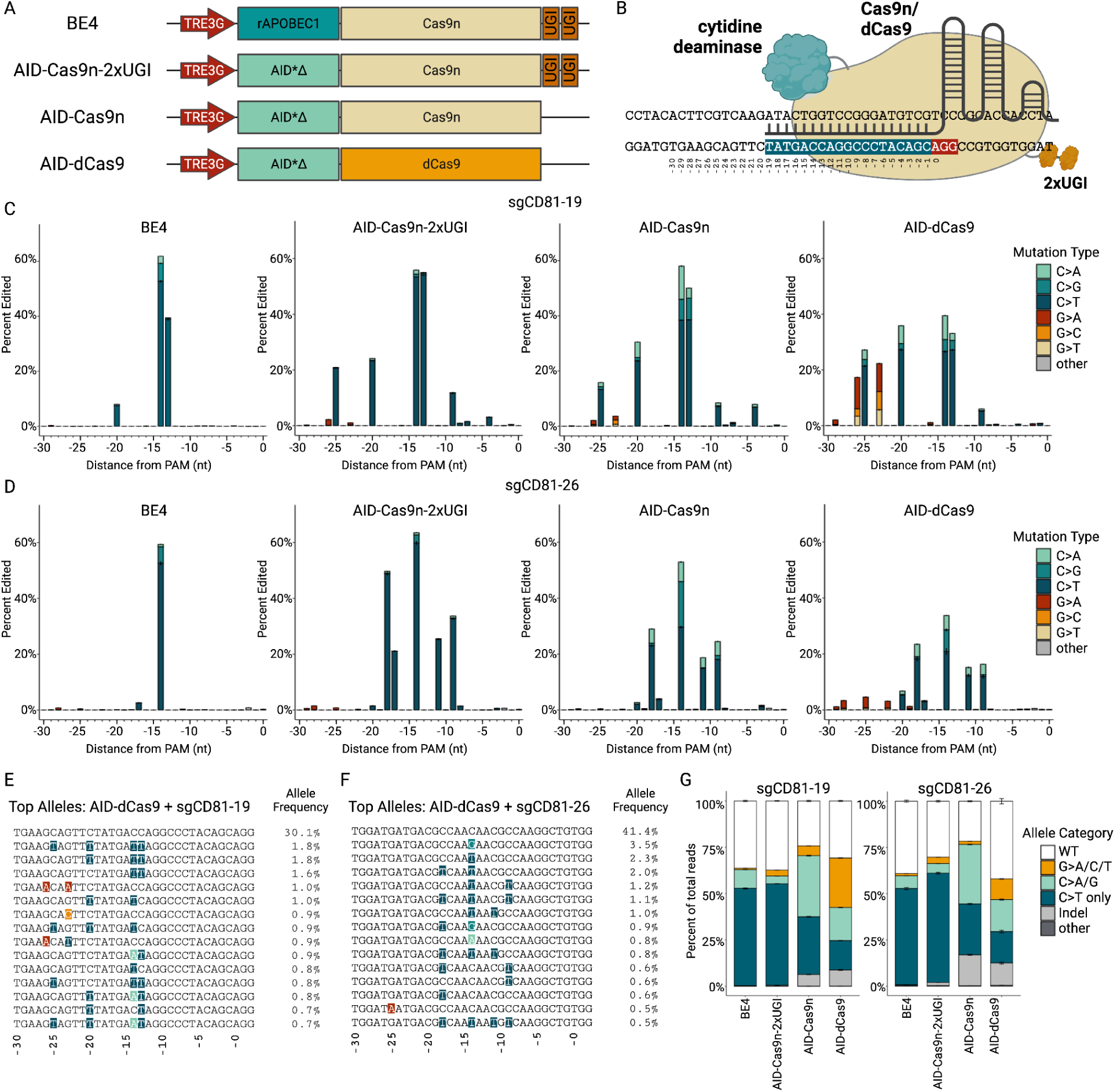
Optimization and characterization of a hypermutagenic base editor, AID-dCas9. **A**. Schematic representation of the base editor domain architectures. All base editors are expressed from the DOX-inducible TRE3G promoter. BE4 contains rat Apobec1, Cas9n (D10A) and 2 UGI domains. All other base editors utilize hyperactive AID, with either Cas9n (D10A) or dCas9, and with or without UGI domains. **B**. Schematic diagram of a base editor with sgRNA bound to a genomic DNA target site. The PAM sequence is highlighted in red, the sgRNA spacer sequence in dark teal. Nucleotides in the editing region are numbered with the 0 indicating the first nucleotide of the PAM sequence. **C-D**. Bar plots showing the percent editing at each nucleotide relative to the PAM for each base editor using sgCD81-19 sgRNA (**C**) or sgCD81-26 sgRNA (**D**). The type of mutation is indicated by color. **E-F**. The top 15 highest frequency alleles with the AID-dCas9 base editor and sgCD81-19 sgRNA (**E)** and sgCD81-26 sgRNA (**F**). **G**. Bar plots showing the percent of total reads in each allele category as indicated by color for each base editor with sgCD81-19 (left) and sgCD81-26 (right) sgRNAs. The C>A/G allele category includes alleles that may contain C>T edits, but do not contain G>A/C/T mutations. For **C-D** and **G**, N = 3 biologically independent replicates. Error bars represent SEM. AID (Activation-induced cytidine deaminase), UGI (Uracil DNA glycosylase inhibitor).

In order to control the timing and duration of genomic exposure to each BE, individual BEs were expressed downstream of a doxycycline (DOX)-inducible promoter within piggyBac plasmids that were stably delivered into PC-9 cells, a model system for mutant EGFR-driven NSCLC (Supplementary Figure 1A). sgRNAs targeting *CD81* (Figure 1B and Supplementary Figure 1A, B) were introduced by lentiviral transduction and editing was assayed by amplicon sequencing of the sgRNA target sites after treatment of the cells with DOX for 6 days. As projected, we observed that the AID-Cas9n-2xUGI base editor had a broader editing window compared to BE4, but retained high editing efficiency and primarily C>T editing specificity (Figure 1C, D and Supplementary Figure 1C, E). Moreover, we found that in the absence of the UGI domains, the editing specificity of AID-Cas9n was no longer restricted to C>T, but also included C>A and C>G edits (Figure 1C, D and Supplementary Figure 1C, E). With AID-dCas9, while the overall editing efficiency was reduced relative to AID-Cas9n, there was a notable increase of G>A, G>C, and G>T edits near the 5’ end of the sgRNA targeting sequence. We interpret this to reflect cytosine editing occurring on the opposite strand (strand bound to the sgRNA) in positions where the DNA is single stranded and accessible to the AID domain (Figure 1C, D and Supplementary Figure 1D, F). Examination of the top 15 alleles generated by AID-dCas9 revealed a diverse array of mutations (Figure 1E, F and Supplementary Figure 1G, H). To determine the degree of allelic diversity attained by each base editor, we assessed the percentage of reads containing C>T edits only, those containing C>A or C>G edits, and those containing G>N edits (Figure 1G and Supplementary Figure 1I). Furthermore, analysis of the total number of alleles and corresponding protein sequences produced by each base editor in combination with each *CD81* sgRNA showed that AID-Cas9n and AID-dCas9 produced on average 5-fold more unique alleles and protein sequences than BE4 (Supplementary Figure 1J, K). Taken together, we find that the AID-dCas9 BE enabled the greatest degree of mutational heterogeneity across the largest window of target base pairs out of all the BE variants tested in our study. Of note, BEs without UGI domains (AID-Cas9n and AID-dCas9) generated a higher incidence of indels. However, with AID-dCas9, these comprise fewer than 10% of the total reads.

**Supplementary Figure 1.**
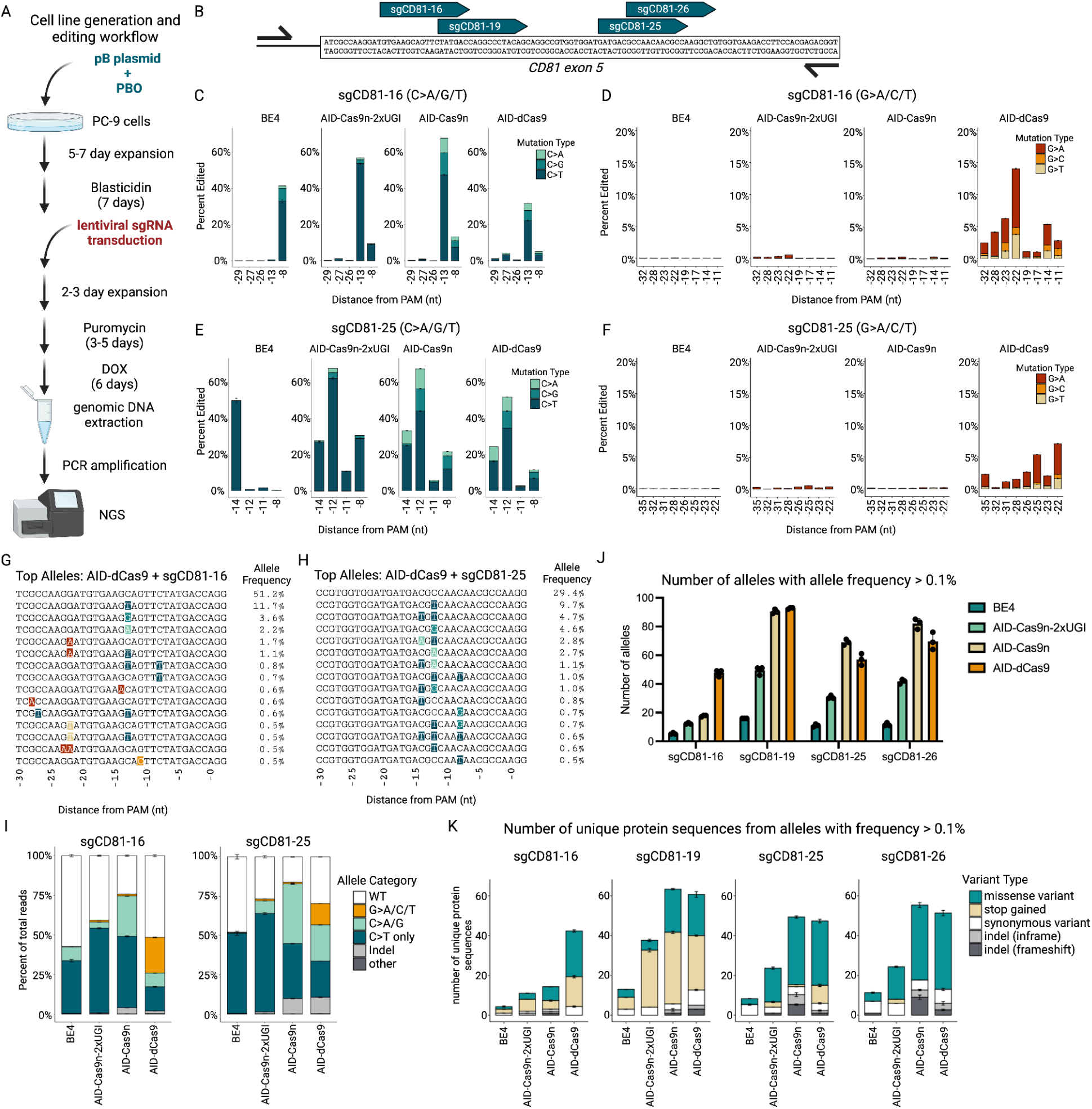
Additional optimization and characterization of the AID-dCas9 base editor. **A**. Overview and timeline of cell line generation and base editing assay. Base editors were expressed from piggyBac plasmids co-expressing a Blasticidin resistance gene, and stably integrated into PC-9 cells after treatment with Blasticidin. Single guide RNAs co-expressed with a Puromycin resistance gene were stably introduced by lentivirus and selected with Puromycin. Cells were then treated with DOX to induce base editor expression for 6 days. Genomic DNA was then isolated, the targeted region amplified and deep sequenced. **B**. Schematic diagram of the *CD81* locus, targeting location of the sgRNAs is indicated by dark teal arrows and the primers used to amplify the region indicated by black arrows. **C**. Bar plots showing the mutation frequency at each C in the editing window, color coded by the type of mutation with sgCD81-16 sgRNA. **D**. Bar plots showing the mutation frequency at each G in the editing window, color coded by type of mutation with sgCD81-16 sgRNA. **E**. Bar plots showing the mutation frequency at each C in the editing window, color coded by the type of mutation with sgCD81-25 sgRNA. **F**. Bar plots showing the mutation frequency at each G in the editing window, color coded by type of mutation with sgCD81-25 sgRNA. **G, H**. Alignment of the top 15 highest frequency alleles edited with the AID-dCas9 base editor and sgCD81-16 (**G**) sgRNA or sgCD81-25 sgRNA (**H**). **I**. Bar plots showing the percent of total reads in each allele category as indicated by color for each base editor with sgCD81-16 (left) and sgCD81-25 (right) sgRNAs. The C>A/G allele category includes alleles that may contain C>T edits, but do not contain G>A/C/T mutations. **J**. Barplot showing the number of alleles with allele frequencies greater than 0.1% with each sgRNA for each base editor as indicated by color. **K**. Barplot showing the number of unique protein sequences encoded by alleles with a frequency of > 0.1% with each variant type indicated by color. For **C-F** and **I-K**, N = 3 independent biological replicates. Error bars represent SEM.

Finally, we confirmed the editing we observed at the *CD81* locus was sgRNA-dependent by expressing a non-targeting control sgRNA (sgNTC) in combination with each BE and assaying the percent editing across the entire *CD81* amplicon (Supplementary Figure 2A). Indeed, across all BEs tested, WT alleles comprised greater than 99% of total reads with an sgNTC (Supplementary Figure 2B).

**Supplementary Figure 2.**
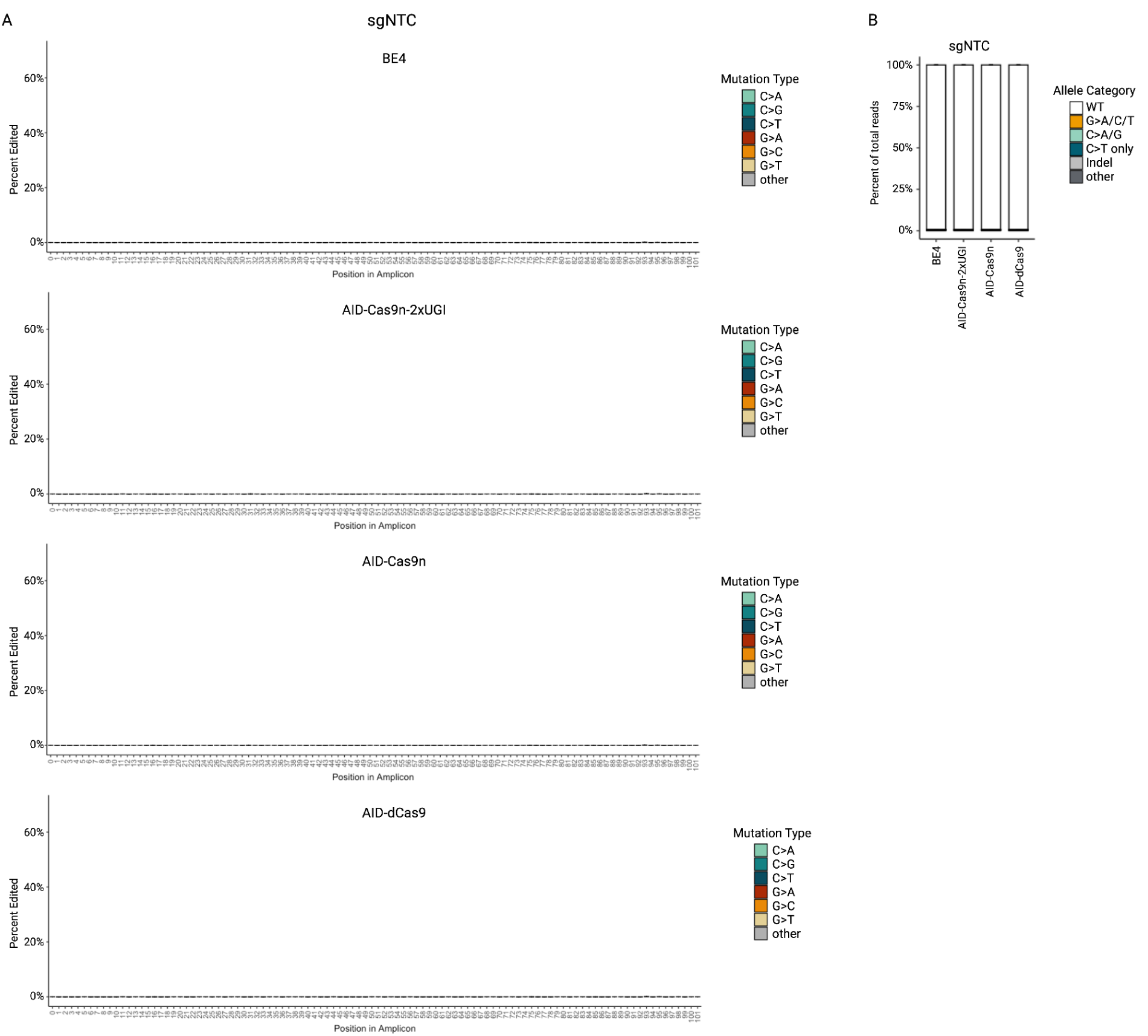
Negligible editing is observed using a non-targeting control (NTC) sgRNA. **A**. Bar plots showing the percent editing at each nucleotide across the CD81 amplicon for each base editor using a non-targeting control (NTC) sgRNA. **B**. Bar plots showing the percent of total reads in each allele category as indicated by color for each base editor using an NTC sgRNA, averaged across all four CD81 targeting sites. N=3 biologically independent replicates, with error bars representing SEM.

We next isolated a high efficiency AID-dCas9-PC-9 clone, based on loss of CD81 expression after induction of AID-dCas9 in the presence of sgCD81-19 (Supplemental Methods). After expanding the clonal population, we characterized the kinetics of base editing by transduction with two distinct *CD81*-specific sgRNAs followed by DOX addition for 0, 1, 3, 6, or 9 days. Cells were then collected and mutagenesis at each locus was analyzed by amplicon-seq (Supplementary Figure 3A, B). Editing was observed as early as 1 day and peaked between days 6 and 9 of DOX treatment, with editing efficiency of up to 60% observed at defined positions within each target site. Both C>N and G>N edits accumulated progressively across the time course, suggesting no significant difference between when C>N edits or G>N edits are catalyzed. Additionally, the number of mutations occurring per allele increased over time, indicating that targets accumulate additional mutations with prolonged exposure to AID-dCas9 (Supplementary Figure 3C). Finally, we observed that the maximum number of unique alleles is achieved between 6 and 9 days of DOX-treatment (Supplementary Figure 3D). As a result of this analysis, we established 6 days of DOX treatment as the ideal duration for base editor expression.

**Supplementary Figure 3.**
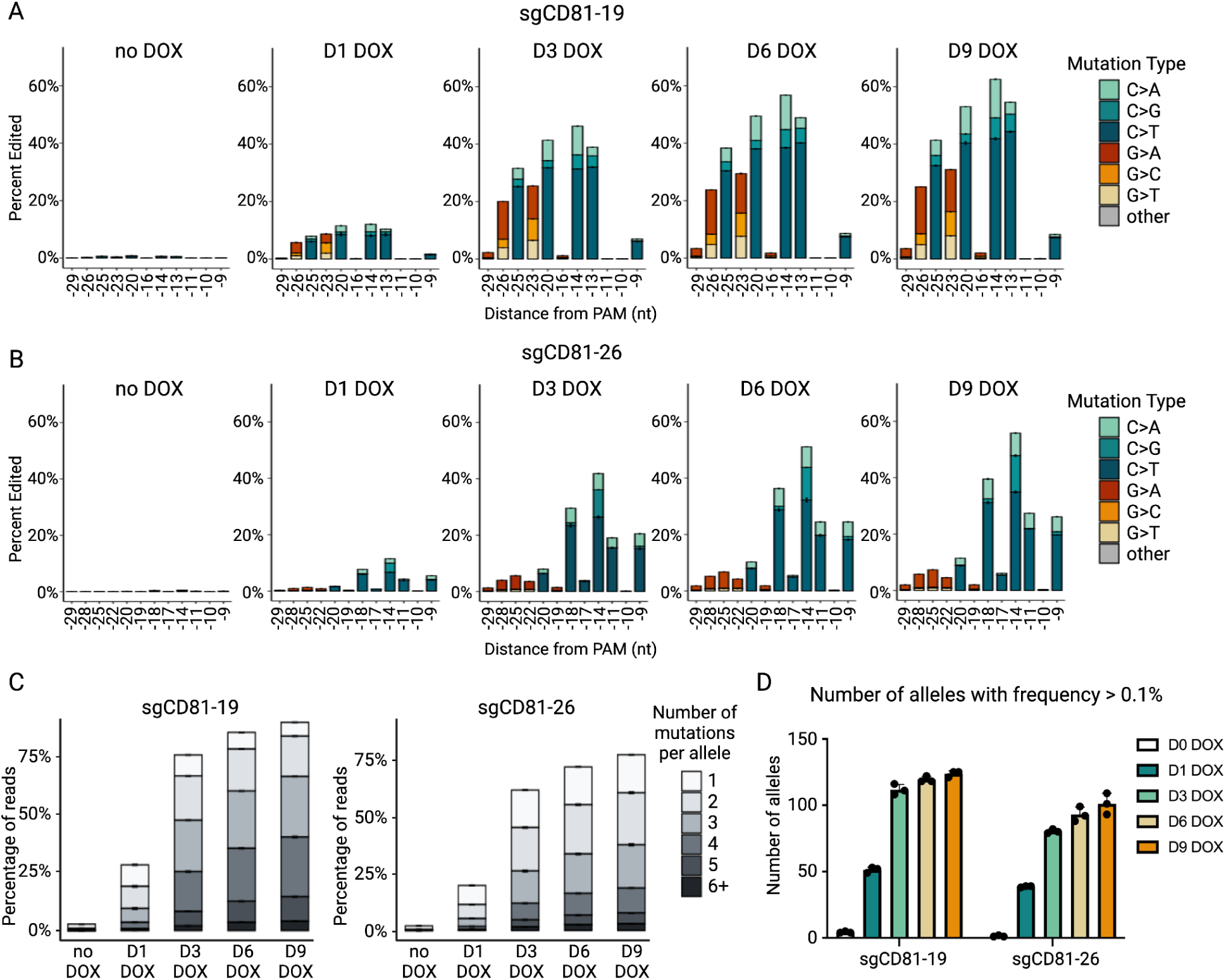
Accumulation of DNA edits over time after DOX-induced AID-dCas9 expression. **A, B.** Barplots showing mutation frequencies of each G and C nucleotide in the editing window across a time course of DOX treatment. The type of mutation is indicated by color, and the sgRNAs used are sgCD81-19 (**A**) sgCD-26 (**B**). **C.** The percentage of reads harboring different numbers of mutations as indicated by the gray scale across a time course of DOX treatment. **D.** Barplot showing the number of alleles with allele frequencies greater than 0.1% with each sgRNA at each DOX time point. For **A-D**, N=3 biologically independent replicates, with error bars representing SEM.

In order to evaluate the degree of saturation that could be obtained in a variant discovery screening context, we applied the AID-dCas9-PC-9 cells described above in a tiled sgRNA assay focused on two distinct loci: a 140 bp region of *CD81* (*CD81*ex5, Figure 2A) and a 216 bp region in the olfactory receptor *OR5M9* (Supplementary Figure 4A). For both target sites, every possible sgRNA with an NGG protospacer-adjacent motif (PAM) was designed, cloned, and stably expressed in an arrayed format, excluding sgRNAs predicted to impact primer binding sites necessary to amplify each region for sequencing analysis. Resulting C>N mutations are shown in teal while G>N mutations displayed in orange (Figure 2A, Supplementary Figure 4A). The base editing profiles from the *CD81*ex5 and *OR5M9* tiling screen data were then used to create a model to predict the editing window for the AID-dCas9 base editor for future sgRNA and screen design. Several sgRNAs were excluded from this analysis (shown in gray text) due to their low DeepHF and CFD scores (53, 54), which correlated with lower base editing activity. This model revealed that while the bulk of C>N edits were localized −21 to −12 bp upstream of the PAM, G>N edits were concentrated beyond the 5’ end of the sgRNA, at −28 to −19 bp upstream of the PAM site (Figure 2B). Using this editing window and a cutoff of 75% maximal editing weight, our model predicted that around 45% of all nucleotides and ∼80% of GC nucleotides in the *CD81* and *OR5M9* amplicons could be edited by AID-dCas9. The observed editing frequencies for these regions, based on direct sequencing, were nearly identical to our predictions (Figure 2C, Supplementary Figure 4B).

**Figure 2.**
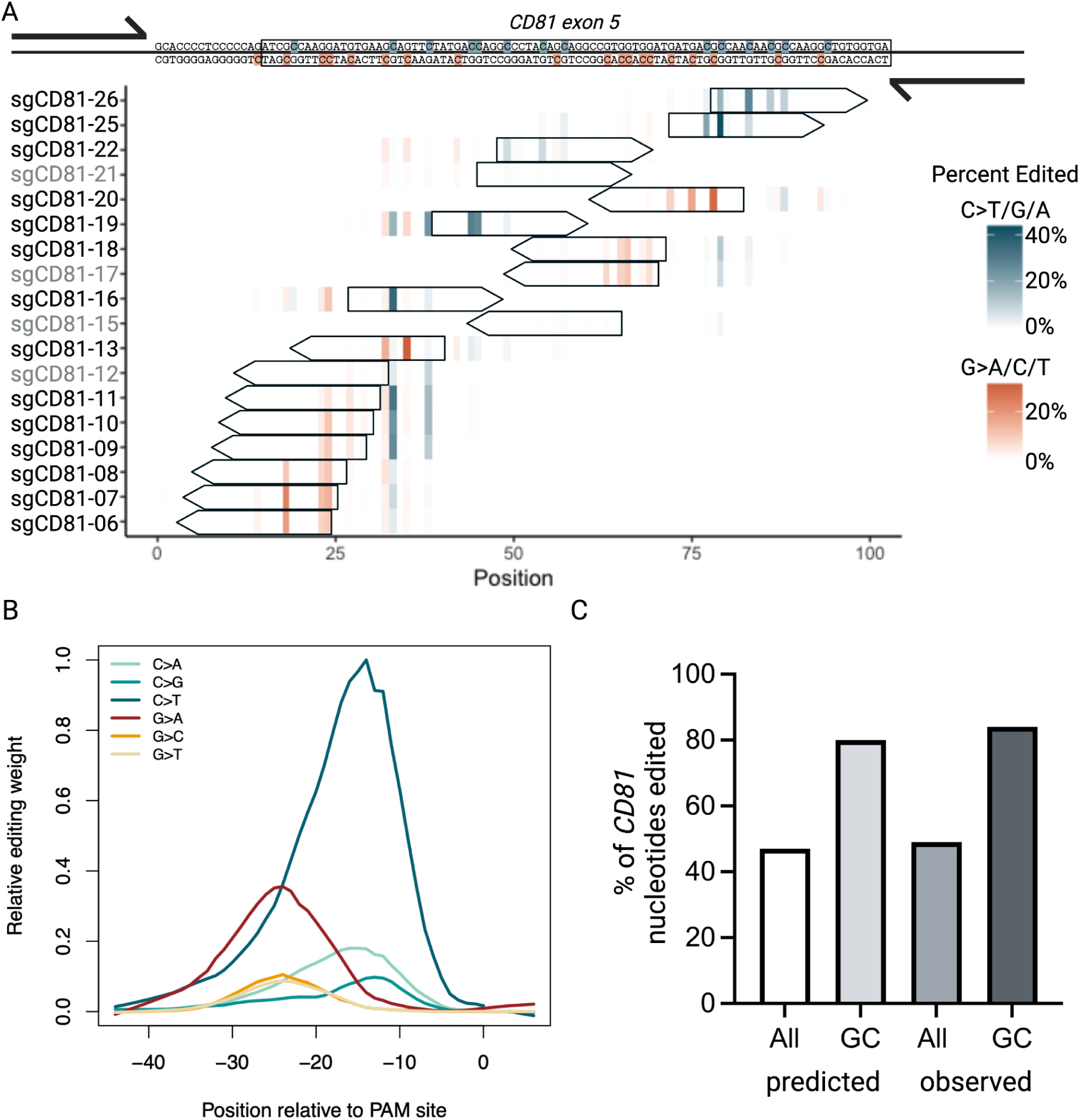
Targeting AID-dCas9 with tiled sgRNAs enables widespread mutagenesis. **A**. Heatmap showing the location and frequency of mutations generated by sgRNAs across a 140 bp amplicon in *CD81* exon 5. Single guide RNAs were individually introduced into a high-efficiency clonal AID-dCas9 PC-9 cell line and assayed for base editing activity. The position of mutations generated by each sgRNA assayed is shown, with the mutation type and mean frequency indicated by the color scale (n=3 biologically independent replicates). The binding location and orientation of each sgRNA is indicated by a rectangular arrow. Targeted Cs and Gs are highlighted in teal and orange, respectively in the DNA sequence shown at the top. sgRNAs with DeepHF and CFD scores both less than 0.5 are indicated by gray text and were excluded from the downstream analysis described in (**B**). **B**. AID-dCas9 editing efficiency as a function of nucleotide position, derived from 36 sgRNAs tiled across a region in *CD81* and *OR5M9*, normalized to the peak editing frequency and lowess smoothed (f=0.3). **C**. Bar graph comparing the percentage of all nucleotides and GC-only nucleotides in the *CD81* exon 5 amplicon which would be predicted to be edited (using all the sgRNAs listed in (**A**) and an editing window corresponding to 75% of maximal for each type of mutation) and the percentage of nucleotides in the *CD81* amplicon that were observed to be edited by the AID-dCas9 base editor.

**Supplementary Figure 4.**
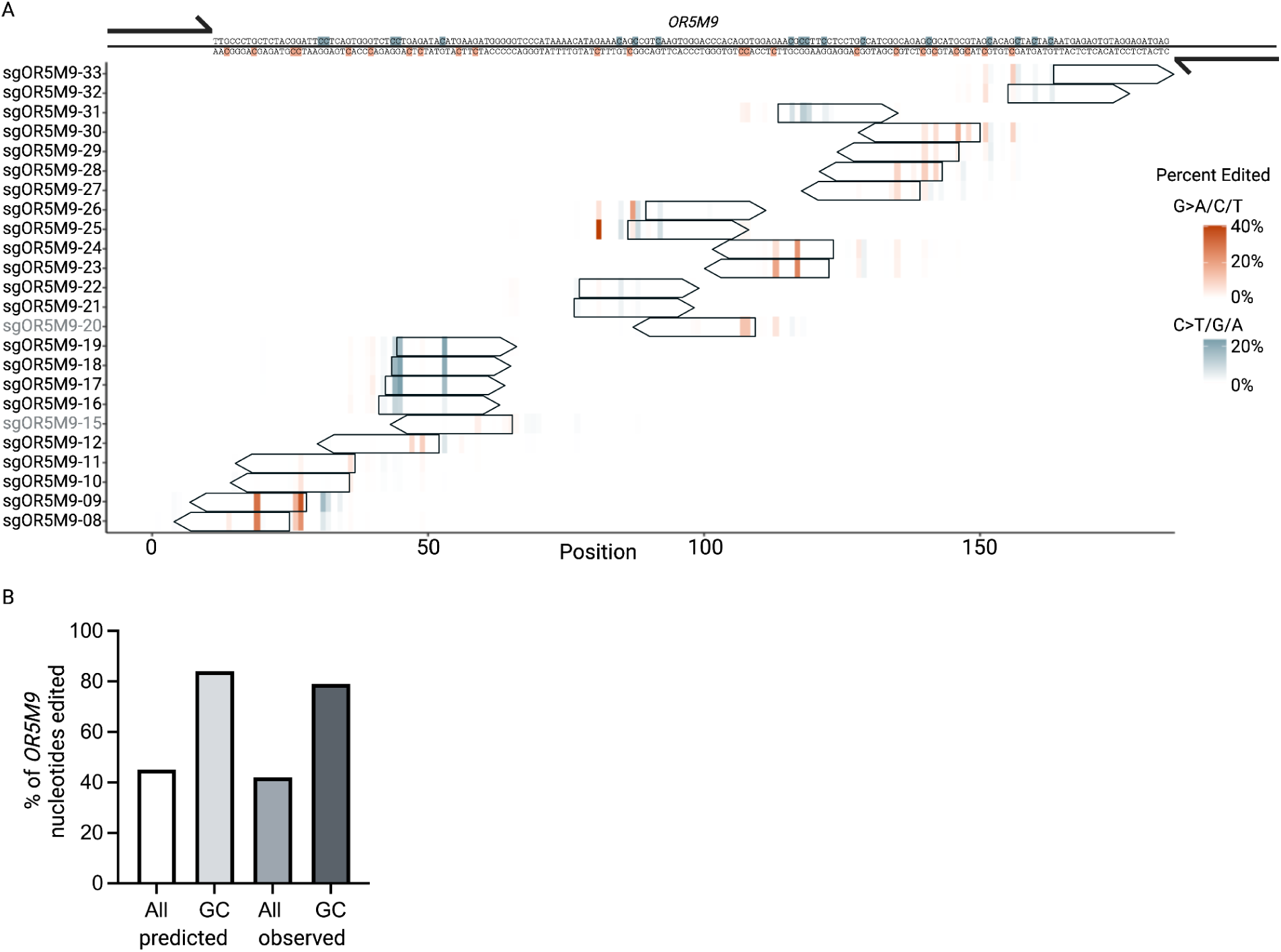
Additional characterization of tiling mutagenesis with AID-dCas9. **A**. Heatmap showing the location and frequency of mutations generated by sgRNAs across a 216 bp amplicon in *OR5M9*. Single guide RNAs were individually introduced into a high-efficiency clonal AID-dCas9 PC-9 cell line and assayed for base editing activity. The position of mutations generated by each sgRNA assayed (n=3 biologically independent replicates) is shown, with the mutation type and mean frequency indicated by the color scale. The binding location and orientation of each sgRNA is indicated by a rectangular arrow. Targeted Cs and Gs are highlighted in teal and orange, respectively in the DNA sequence shown at the top. sgRNAs with DeepHF and CFD scores both less than 0.5 are indicated by gray text and were excluded from the editing weights calculation (**Figure 2B**). **B**. Bar graph comparing the percentage of all nucleotides and GC-only nucleotides in the *OR5M9* amplicon which would be predicted to be edited (using all the sgRNAs listed in (**A**) and an editing window corresponding to 75% of maximal for each type of mutation) and the percentage of nucleotides in the *OR5M9* amplicon that were observed to be edited by the AID-dCas9 base editor.

After demonstrating that AID-dCas9 was capable of generating both a high density and variety of mutations across a given locus, we wanted to further assess its ability to generate functional variants in a bona fide screening context. Therefore, we established a series of proof-of-concept screens and subsequent validation studies based on the paradigm of EGFR inhibitor resistance in NSCLC. First generation EGFR inhibitors, such as erlotinib, dampen elevated signaling of the mutant receptor by competitively blocking ATP binding to the kinase domain of EGFR. The EGFR T790M mutation enables resistance to this class of small molecules by preventing their binding. Osimertinib is a third generation EGFR inhibitor which forms a covalent interaction with C797, and while maintaining potent activity towards the common EGFR mutants, is also able to potently suppress T790M containing variants (6) (Figure 3A). As might be expected for a covalent molecule, the most frequently reported on-target (*cis*) osimertinib resistance mutation is C797X. However, the overall frequency of on-target mutations is much less frequent than with first generation molecules. With osimertinib, the clinical resistance landscape includes SNV variants in the downstream mitogen-activating protein kinase (MAPK) signaling pathway such as *BRAF*, amplification of related receptors such as *MET* or *HER2*, alterations in cell cycle genes, and as-yet-unexplained mechanisms (by far the largest class representing 30-50% of resistant samples) (55).

**Figure 3.**
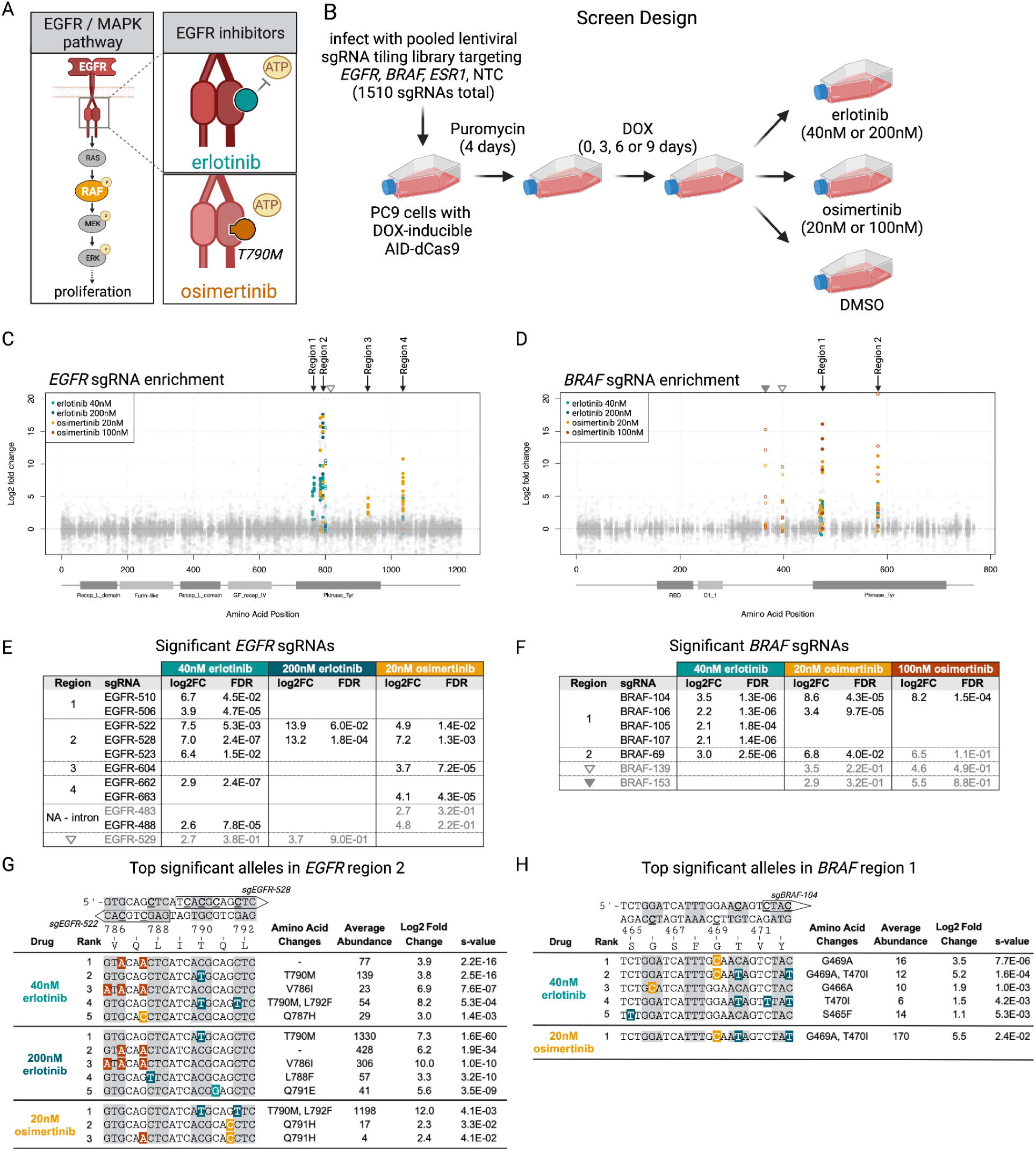
Pooled base editing screen for EGFR inhibitor resistance. **A**. Schematic diagram of the signaling pathway downstream of EGFR and mechanism of action of the EGFR inhibitors, erlotinib and osimertinib. Erlotinib competes with ATP for binding to the ATP binding pocket, while osimertinib preferentially and irreversibly binds to EGFR harboring T790M or other activating mutations. **B.** Schematic overview of the base editing screen design. A pooled lentiviral library containing sgRNAs tiling the coding regions of *EGFR*, *BRAF* and *ESR1* was transduced into a high-efficiency clonal DOX-inducible AID-dCas9 PC-9 cell line, followed by Puromycin selection and DOX treatment for either 0, 3, 6 or 9 days. Cells were then treated with 40nM or 200nM doses of erlotinib or 20nM or 100nM doses of osimertinib. **C**, **D.** Scatter plot showing the log2 fold change relative to the DMSO controls of significantly enriched sgRNAs in *EGFR* (**C**) and *BRAF* (**D**) from samples treated with doses of erlotinib and osimertinib as indicated by color. Significance was determined by log2 fold change > 2 and FDR < 0.05, treating the day 3, 6 and 9 samples as replicates. Open circles indicate the location and log2 fold change of sgRNAs which we consider to be weak hits (log2 fold change >2, FDR > 0.05). Significant sgRNAs cluster in four regions in *EGFR* and two regions in *BRAF* and the location of weak hits are indicated by gray open or filled triangles. **E, F.** List of significantly enriched and weak hit sgRNAs in *EGFR* (**E**) and (**F**) *BRAF* grouped by region and separated into columns which indicate the dose of erlotinib or osimertinib with which the cells were treated. Gray text indicates the weak hit sgRNAs. Blank spaces indicate the sgRNA was not significantly enriched in that condition. **G, H.** The top 5 alleles ranked by s-value for each drug condition in *EGFR* region 2 (**G**) or *BRAF* region 1 (**H**). The location of the relevant sgRNAs targeting the region are indicated by rectangular arrows overlapping the double stranded DNA sequence, with each codon highlighted by an alternating gray and white background and edited cytosines underlined. The mutated alleles are shown with mutations highlighted in red/orange/cream for G-to-N edits and shades of teal for T-to-N edits. The resulting amino acid changes, relative abundance of the allele in the population, log2 fold change and s-value are reported for each allele. The significance cutoff was s-value <0.05, log2 fold change >1. In cases where there were less than 5 significant alleles, all significant alleles are shown. The full list of significant variants can be found in Supplementary Data.

Owing to the potential for discovering novel resistance variants in *cis* (drug target) or *trans* (secondary locus), we devised a screen to select for mutations in EGFR or BRAF, a downstream effector within the EGFR/MAPK pathway (Figure 3A), that reduce sensitivity to EGFR inhibitors. Here, a pool of sgRNAs was designed to tile all possible NGG-containing target sites within the coding regions of *EGFR* and *BRAF* (Supplementary Figure 5A, B). Also included were sgRNAs tiled across a non-MAPK pathway gene, *ESR1*, and 50 non-targeting sgRNA controls, both serving as negative controls. sgRNAs with the potential to bind many sites in the genome or with sequences that could interfere with library cloning were excluded. Altogether, the *EGFR* and *BRAF* sgRNAs were predicted to edit approximately 40% and 30% of all coding bases, and 72% and 64% of GC base pairs, respectively (Supplementary Figure 5C). The AID-dCas9-PC-9 clonal line was chosen for the screen, due to the potent antiproliferative activity of EGFR inhibitors in these cells. Importantly, genotyping confirmed the AID-dCas9-PC-9 clone as wild type for position T790, and based on our predictive model, several sgRNAs within our library had the potential for generating the T790M variant. Prior to delivery of the sgRNA library into AID-dCas9-PC-9 cells, we performed dose response analyses for both erlotinib and osimertinib, which demonstrated equivalent sensitivity between the AID-dCas9-PC-9 and parental PC-9s across conditions (Supplementary Figure 5D, E). Subsequently, for the erlotinib arm, we chose a low dose (40nM, IC80) to select for a broad range of resistant clones and a high dose (200nM, >IC95) more comparable to the serum concentrations achieved in patients (56) to select for more robust and potentially clinically relevant variants. For the osimertinib arm, we prioritized two doses (20nM, IC95 and 100nM, >IC95), which are inline with osimertinib concentrations detected in patient blood samples following standard dosing (57, 58) to enrich for stronger, more pertinent resistance variants.

**Supplementary Figure 5.**
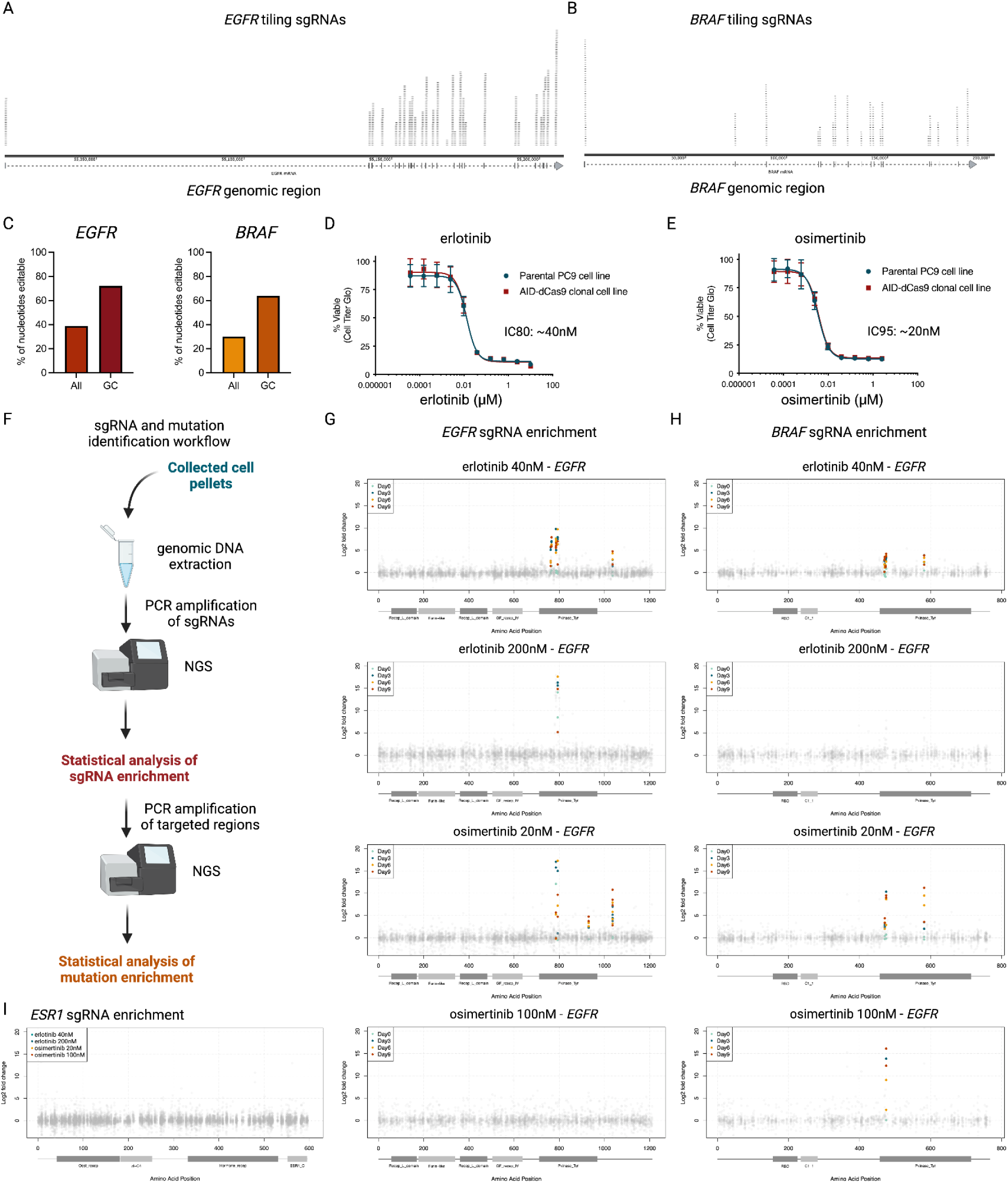
Additional information on the tiling screen for EGFR inhibitor resistance mutations. **A, B**. Schematic of the sgRNA distribution across the *EGFR* (**A**) and *BRAF* (**B**) genomic region. **C**. Bar graphs indicating the percentage of all nucleotides and CG nucleotides only in the coding region of *EGFR* (left) and *BRAF* (right) that predicted to be editable using the AID-dCas9 base editor (see **Figure 2B** for the editing window and weights, calculated down to 75%-maximal editing activity for each type of mutation). **D, E**. Dose response curves showing the viability of parental and engineered AID-dCas9 clonal PC-9 cell lines treated with escalating doses of erlotinib (**B**) and osimertinib (**C**). **F**. Schematic diagram of the sgRNA and mutation identification workflow. **G, H**. Significantly enriched over DMSO control (log2 fold change >2, FDR < 0.05) *EGFR* (**G**) and *BRAF* (**H**) sgRNAs at each drug and dose color coded by the number of days each sample was treated with doxycycline to induce base editor expression. **I**. Enrichment of *ESR1* sgRNAs in the EGFR inhibitor screen (none are significant).

Following lentiviral transduction of the sgRNA library at low MOI (0.3), AID-dCas9-PC-9 cells were treated with a short time course of DOX to induce base editing and collect the broadest range of variants possible (Figure 3B). DOX was removed from the cultures, and each cell population was exposed to either erlotinib or osimertinib, using the conditions described above, or DMSO (negative control). Surviving cells were collected after expansion and processed to identify the sgRNAs from each condition (Supplementary Figure 5F). We did not see noticeable differences in sgRNA enrichment between the 3, 6 and 9 day DOX time points (Supplementary Figure 5G, H), so we treated these samples as replicates to increase the statistical power of the screen. Sequence analysis of sgRNAs from cells collected across all conditions, relative to DMSO, revealed four regions in *EGFR* and two in *BRAF* enriched with highly significant sgRNA enrichment (Figure 3C, D). One additional *EGFR* sgRNA adjacent to region 2 and two *BRAF* sgRNAs had high enrichment (log2 fold change) across several drug conditions, but did not meet the significance threshold (FDR < 0.05), so were designated as weak hits (open circles). No sgRNAs were determined to be significantly enriched in the *ESR1* control set (Supplementary Figure 5I). Significant sgRNAs overlapping Region 2 of *EGFR*, which mapped to T790, were prevalent in all but the high dose osimertinib arm (Figure 3E), while Region 1 in *BRAF* was enriched in all but the high dose erlotinib arm (Figure 3F). Targeted genomic sequencing of cells collected from both erlotinib and the low dose osimertinib arms revealed that mutations resulting in the T790M variant were indeed present and at high proportion relative to other alleles at that locus (Figure 3G). Notably, in the lower dose arms for both erlotinib and osimertinib, we observed T790M in *cis* with L792F, a recently described mutation conferring resistance to third generation inhibitors (59–62). Variants at positions V786 and L788 were consistent with rare alleles observed in lung cancer patients (63, 64), while the Q791H allele had been found at low frequency in osimertinib-resistant tumors (60). We also observed enrichment of silent mutations in erlotinib, which may represent high efficiency ‘hitchhiking’ mutations made by sgEGFR-522 alongside T790M on an alternate allele. In parallel, sequencing of the *BRAF* region 1 locus, which was enriched in 3 of the drug treatments, revealed BRAF G469A-producing mutations, along with T470I variants on their own or in *cis* with G469A, as top hits for the lower dose erlotinib and osimertinib arms (Figure 3H). BRAF G469A mutations have recently been reported to confer resistance to osimertinib (65). We also sequenced *EGFR* region 1 from the low dose erlotinib condition and identified EGFR S768I, a clinically-relevant variant observed in patients (66) (Supplementary Data). While no statistically significant variants emerged from sequencing of *BRAF* region 2, this regions maps within the protein kinase domain, with proximity to E586-D594; a known hot-spot for mutations in cancer patients or patients with reduced sensitivity to inhibitors of varying MAPK pathway effectors (67, 68), albeit without links to EGFR inhibitor resistance. The ability of our screen to recover known variants, including complex alleles containing multiple amino acid changes, provided validation that the AID-dCas9 system can create diverse and functional mutations.

For granular accounting of variants that enabled growth in the presence of erlotinib or osimertinib after editing with AID-dCas9 we performed a secondary screen using select sgRNAs identified in the primary screen coupled to a single cell genotyping readout (Figure 4A). Single cell (sc) genotyping was performed using the Tapestri platform, which has been applied previously to interrogate genome editing outcomes in a screening context (29, 69, 70). Vectors expressing the most significantly enriched sgRNAs for *EGFR* regions 1-4 and *BRAF* regions 1-2 were pooled, along with the weak hit sgRNAs. After delivery of this pool into AID-dCas9-PC-9 cells and induction of editing for 6 days, the cells were selected using doses corresponding to IC95 for both erlotinib (100nM) and osimertinib (20nM); dosages chosen to enrich for potent modifiers of the drug response (Figure 4A). Loci targeted by each sgRNA were then PCR amplified/indexed and sequenced. Approximately 4, 000 cells were recovered per sample, with read coverage varying by amplicon from approximately 20, 000 to 200, 000 reads per sample per amplicon (Figure 4B, C). We applied filtering criteria to reduce noise in the data and applied a Bayesian negative binomial regression model to identify enriched variants (see Supplemental Methods). We first examined significantly-enriched variants in *EGFR* (s-value < 0.01). The most prevalent amino acid changes occured upon erlotinib treatment and clustered in the EGFR Kinase Domain, including amino acid T790, with T790M and adjacent Q791E mutations being the most enriched (Figure 4D). L792F and other T790 proximal mutations were also significantly enriched in erlotinib. To examine the allelic context of these mutations, we aggregated the single cell data and performed a pseudo bulk analysis. The top three significantly-enriched alleles ranked by s-value in *EGFR* with erlotinib all include a T790M mutation (T790M, T790M/Q791E and T790M/L792F), supporting the notion that this mutation is a dominant force of resistance to erlotinib (Figure 4E). However, in osimertinib, the only significantly enriched allele is T790M/L792F, consistent with the observation that additional mutations are required in *cis* with T790M (ie. L792F) to confer resistance to osimertinib (Figure 4F) (9).

**Figure 4.**
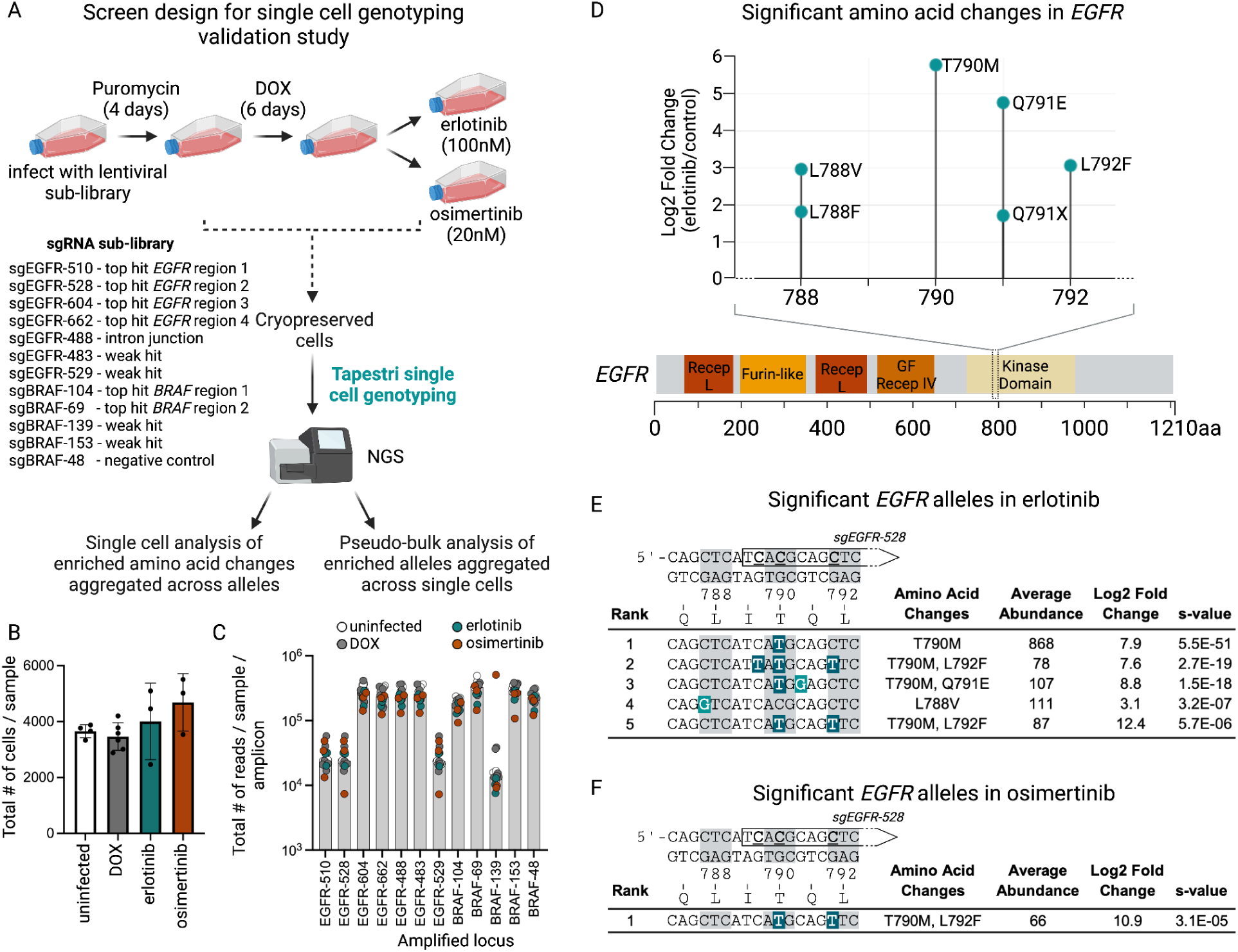
Single cell genotyping of mutations in EGFR conferring resistance to EGFR inhibitors. **A**. Schematic diagram of the workflow of the single cell genotyping screen performed with a sub-library consisting of the top enriched sgRNAs for each region in *EGFR* and *BRAF*, several weaker hits and a negative control sgRNA. Base editor expressing cells were infected with the sub-library, treated with Puromycin, followed by doxycycline and finally an IC95 dose of either erlotinib or osimertinib. Cell pellets were cryopreserved before performing the Tapestri single cell genotyping workflow. Analysis of enriched mutations was performed on the single cells data, while analysis of enriched alleles was performed on pseudo-bulk data from the same experiment. **B**. Bar graphs showing the number of cells obtained from each sample (uninfected, n=4; DOX, n=6; erlotinib, n=3; osimertinib, n=3). **C**. The total number of reads in each sample per amplicon. **D**. Lollipop plot showing the log2 fold change in drug treatment over doxycycline control of amino acid changes in erlotinib (note: no significant mutations were identified in *EGFR* in osimertinib) and where along the *EGFR* gene they are located. **E, F**. Tables showing the enrichment of the top 5 *EGFR* alleles identified in erlotinib (**E**) and osimertinib (**F**) ranked by s-value. The location of the relevant sgRNA targeting the region is indicated by a rectangular arrow overlapping the double stranded DNA sequence, with each codon highlighted by an alternating gray and white background and edited cytosines underlined. The mutated alleles are shown with mutations highlighted in red/orange/cream for G-to-N edits and shades of teal for T-to-N edits. The resulting amino acid changes, relative abundance of the allele in the population, log2 fold change and s-value are reported for each allele. The significance cutoff was s-value <0.001, log2 fold change >1. In cases where there were less than 5 significant alleles, all significant alleles are shown. The full list of significant alleles is included in Supplementary Data.

Performing a similar single cell analysis of individual amino acid changes in BRAF revealed three regions with enriched variants in osimertinib (Figure 5A). The most substantial of these were G469 mutations, several of which have been previously reported to confer resistance to osimertinib (65). Another cluster of enriched mutations were located at G593/D594 within the BRAF kinase domain. Finally, mutations adjacent to S365, a phosphorylated residue that serves as a binding site for inhibitory 14-3-3 proteins, were also identified. To understand the behavior of these mutations at the allelic level, we proceeded to examine the enrichment of alleles at each of these regions in pseudo bulk. This revealed G469A as the major mutation in the BRAF P-loop region, either alone or in combination with T470I or T470K (Figure 5B). Notably, alleles with mutations in E586K and D594N in *cis* were strongly enriched in osimertinib, approximately an order of magnitude greater than E586K alone (Figure 5C). Both of these mutations have been described independently in tumors (71), although their role in driving resistance to osimertinib has not previously been shown. Lastly, two alleles encoding A366P mutations were significantly enriched in osimertinib (Figure 5D). While this variant has been observed in melanoma samples, it was assumed to be a passenger mutation (72). In contrast to osimertinib, no individual *BRAF* mutations were identified as significantly enriched in erlotinib, and only one *BRAF* allele encoding a G469A and T470K mutation in *cis* were significantly enriched in erlotinib (Figure 5E).

**Figure 5.**
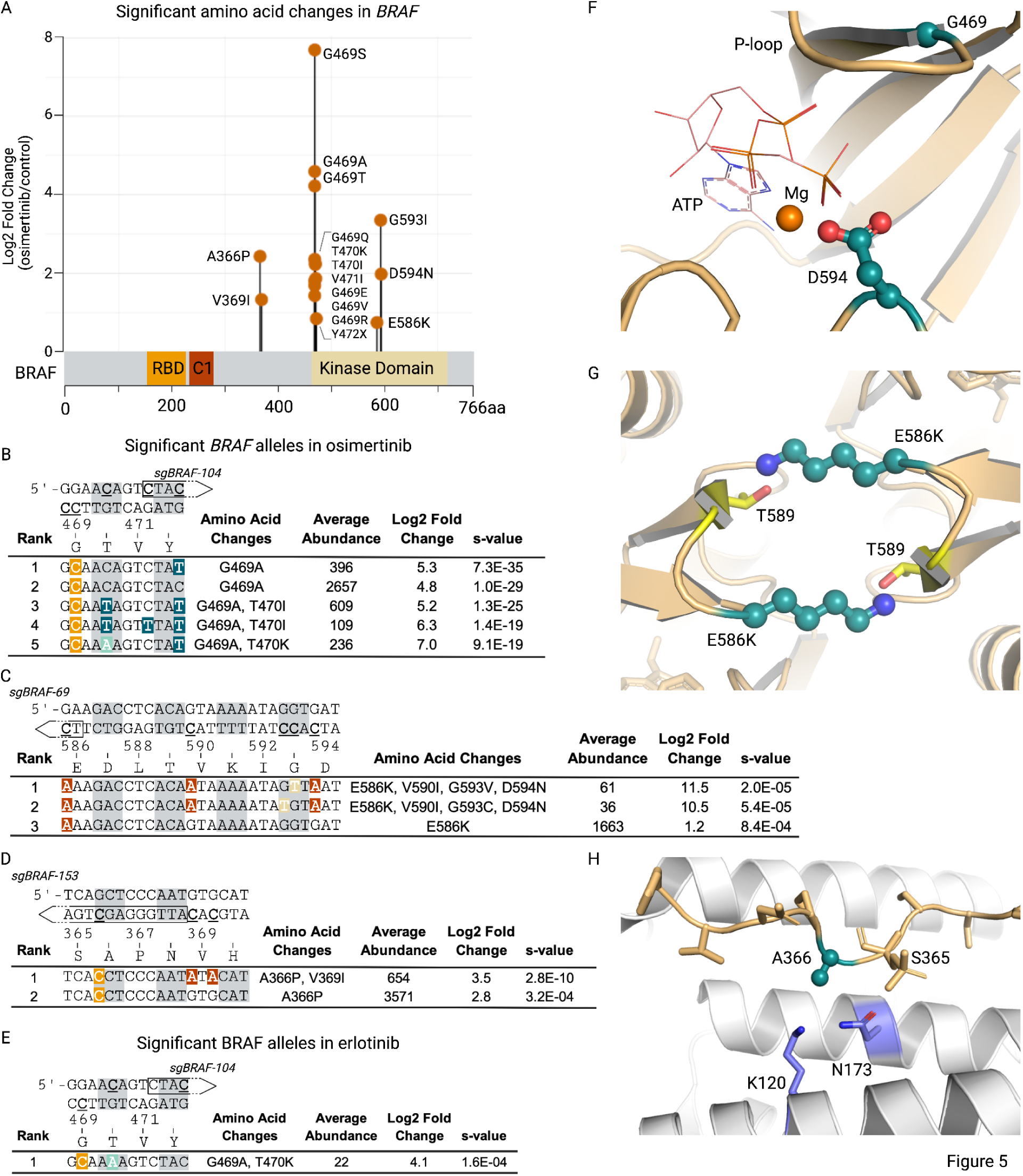
Single cell genotyping data of mutations in *BRAF* enriched following treatment with EGFR inhibitors. **A.** Lollipop plot showing the log2 fold change in drug treatment over doxycycline control of amino acid changes in osimertinib (note: no significant mutations were identified in *BRAF* in erlotinib) and where along the *BRAF* gene they are located. **B-E.** Tables showing the enrichment of the top 5 *BRAF* alleles identified in osimertinib (**B-D**) and erlotinib (**E**) ranked by s-value. The location of the relevant sgRNA targeting the region is indicated by a rectangular arrow overlapping the double stranded DNA sequence, with each codon highlighted by an alternating gray and white background and edited cytosines underlined. The mutated alleles are shown with mutations highlighted in red/orange/cream for G-to-N edits and shades of teal for T-to-N edits. The resulting amino acid changes, relative abundance of the allele in the population, log2 fold change and s-value are reported for each allele. The significance cutoff was s-value <0.001, log2 fold change >1. In cases where there were less than 5 significant alleles, all significant alleles are shown. The full list of significant alleles is included in Supplemental Data. **F.** Structure of the ATP-binding pocket of BRAF showing the location of the mutated amino acids implicated in resistance to osimertinib, G469 in the P-loop and D594 in the DFG loop (PDB code 6U2G). **G.** Modeling of the impact of an E586K mutation on the BRAF dimer interface (PDB code 6XFP), potentially stabilizing a hydrogen bonding interaction of E586K on one protomer with T589 in the second protomer in the partner molecule and vice versa, further promoting dimerization. **H.** Structure showing the location of BRAF A366 at the interface between BRAF and 14-3-3 proteins (PDB code 6NYB), including phosphorylated S365 (pS365) in BRAF. The A366P mutation in BRAF would sterically clash with K120 and N173 in 14-3-3 preventing pS365 from binding to 14-3-3 and stabilizing the inactive conformation of BRAF.

To understand the potential impact of these variants on BRAF activity, we used available structural and associated biochemical data to visualize the location of the mutated amino acids and predict the functional consequences. BRAF is active when dimerized with other RAF (ARAF, BRAF or CRAF) molecules, thus mutations that promote BRAF dimerization would be predicted to result in increased BRAF kinase activity, which could override the pathway blockade caused by EGFR inhibition (73–75). ATP has a negative regulatory effect on BRAF activity by stabilizing an inactive monomeric conformation of BRAF kinase domain in the absence of 14-3-3 induced dimerization (76, 77). G469 in the P-loop (residues 462-469) and D594 in the DFG motif (residues 594-596) in BRAF form a part of the ATP binding site (Figure 5F). Substitution of glycine for alanine at 469 would affect the conformation of the P-loop resulting in a less flexible backbone due to the larger sidechain, making it insensitive to ATP-mediated negative regulation, and prone to activation *in trans* of another BRAF molecule through dimerization. D594N, eliminates the negative charge and would be expected to disrupt the coordination of the Mg ion with ATP, and similar to the G469A mutation, a D594N mutation would also relieve the negative regulatory effects on ATP, promote RAF dimers and result in RAF activation. In addition, modeling of the E586K mutation into the BRAF dimer structure (PDB code 6CFP) shows a potential mechanism by which E568K promotes dimerization through interactions with T589 on the opposite BRAF partner molecule and vice versa (Figure 5G). Thus it is reasonable to conclude that the combined effect of E586K and D594N would result in a strong propensity to form BRAF dimers, leading to increased BRAF activity and MAPK pathway signaling. Finally, we examined the interface between BRAF and 14-3-3 where the phosphorylated S365 (pS365), a known inhibitory phosphorylation site, is located (78). An adjacent A366P substitution would be expected to sterically clash with K120 and N173 residues in 14-3-3, preventing binding of the pS365 containing region of BRAF to 14-3-3, and relieving the negative regulatory effect of 14-3-3 at this site, resulting in RAF dimers and pathway activation (Figure 5H).

Encouraged by the results of our in situ validation, we sought to characterize the effects of single or multi-amino acid changes using complementary over-expression assays, with emphasis on the newly-identified *BRAF* alleles. For this, we engineered a series of mutations into DOX-inducible *EGFR* and *BRAF* transgenes, and stably delivered these to parental PC-9 cells (Figure 6A, Supplemental Figure 6A). *EGFR* transgenes, incorporating the activating exon 19 deletion (del746-750) present in PC-9 cells, with or without T790M and T790M/L792F, served as positive controls for our assay. Transgene-derived *EGFR* expression alone improved survival in the presence of both drugs. However, viability was yet-further enhanced across a range of doses with T790M (erlotinib) and T790M/L792F (erlotinib and osimertinib) (Figure 6B). Similarly, a marked improvement in survival across drugs and doses was observed following over-expression of *BRAF*. While BRAF G469A was able to provide resistance to both erlotinib and osimertinib, the presence of T470I or T470K in *cis* with G469A did not enhance this effect, suggesting that mutations at T470 were potentially bystanders and a result of localized, highly-efficient editing by AID-dCas9 (Figure 6B). By contrast, the presence of D594N, G593V, and V590I alongside E586K (quadruple mutant) substantially improved cell viability over E586K alone across a dose response of either erlotinib or osimertinib (Figure 6B). Finally, we also observed the presence of an A366P mutation in BRAF enhanced viability, although to a modest extent, consistent with the weaker effects observed with this variant in our primary and secondary BE screens (Figure 5B). Expression of BRAF single (G469A, A366P) and compound (E586K/V590I/G593V/D594N [KIVN]) variants led to consistent upregulation of phosphorylated ERK and MEK in the presence of erlotinib and osimertinib, relative to controls, confirming that these variants rescued signaling through the MAPK pathway (Figure 6C). To assess the generalizability of these mutations for conferring drug resistance, we introduced the *EGFR* and *BRAF* mutant transgenes into a separate EGFR mutant and highly EGFR inhibitor responsive NSCLC cell line, HCC827. Dose response analysis with osimertinib resulted in similar increases in viability for each of the mutations tested (Supplementary Fig 6B). Interestingly, in this cellular context, the BRAF T470I or T470K variants in *cis* with G469A provided a modest improvement in viability.

**Figure 6.**
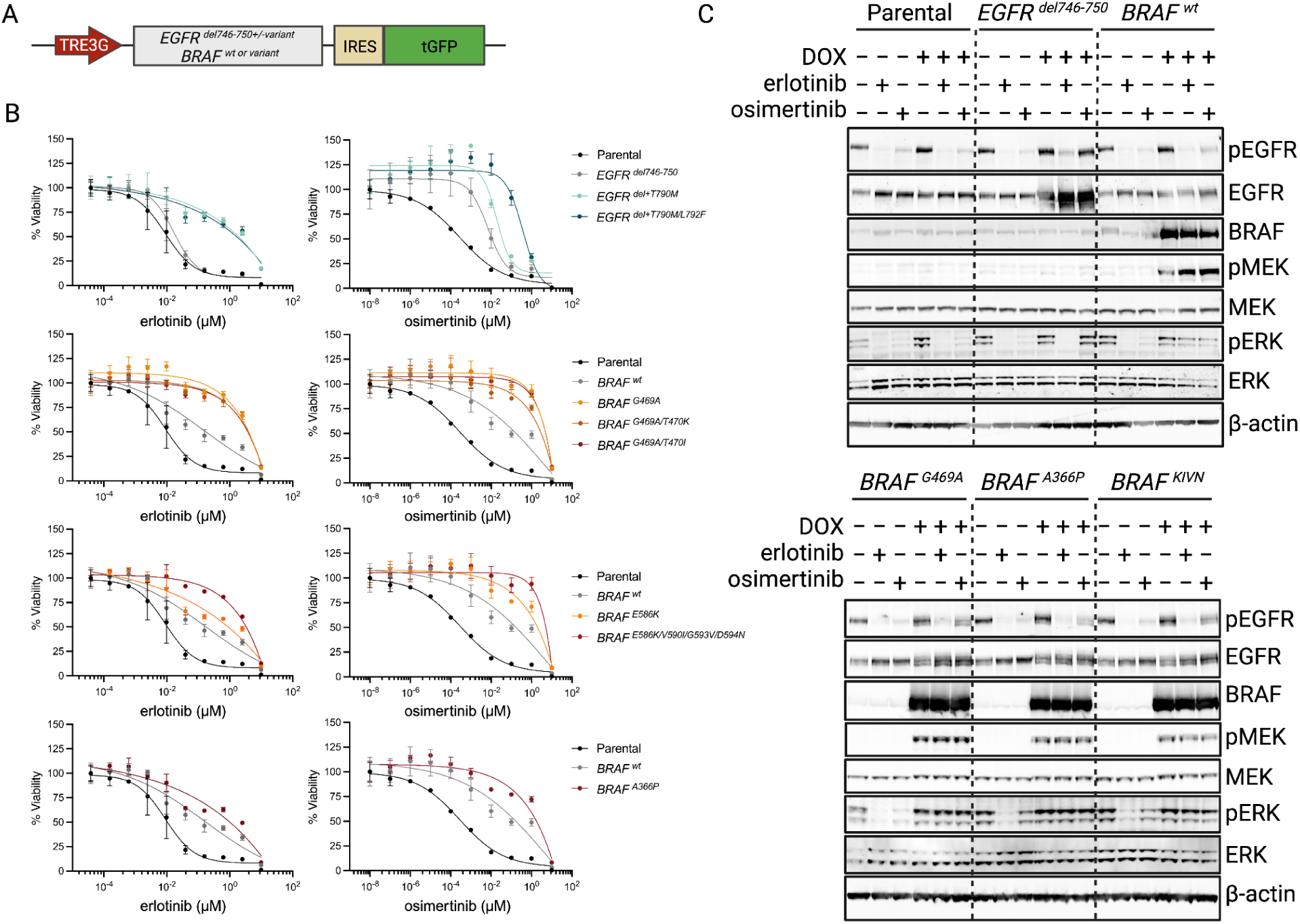
Expression of DOX-inducible *EGFR* and *BRAF* mutant transgenes recapitulates the EGFR inhibitor resistance phenotype. **A.** Schematic representation of the *EGFR* or *BRAF* doxycycline-inducible transgenes. EGFR mutations were introduced into the *EGFR* del746-750 transgene context to mimic as closely as possible the allele created in the PC-9 cells. **B.** Dose-response curves showing the viability of PC-9 cells expressing the specified *EGFR* or *BRAF* transgenes at escalating erlotinib (left) or osimertinib (right) concentrations. **C.** Western blot analysis for markers of MAPK pathway activation, including phospho-EGFR, phospho-ERK and phospho-MEK in cell lines harboring doxycycline-inducible mutant *EGFR* or *BRAF* transgenes, in the presence or absence of DOX, erlotinib, osimertinib.

**Supplementary Figure 6.**
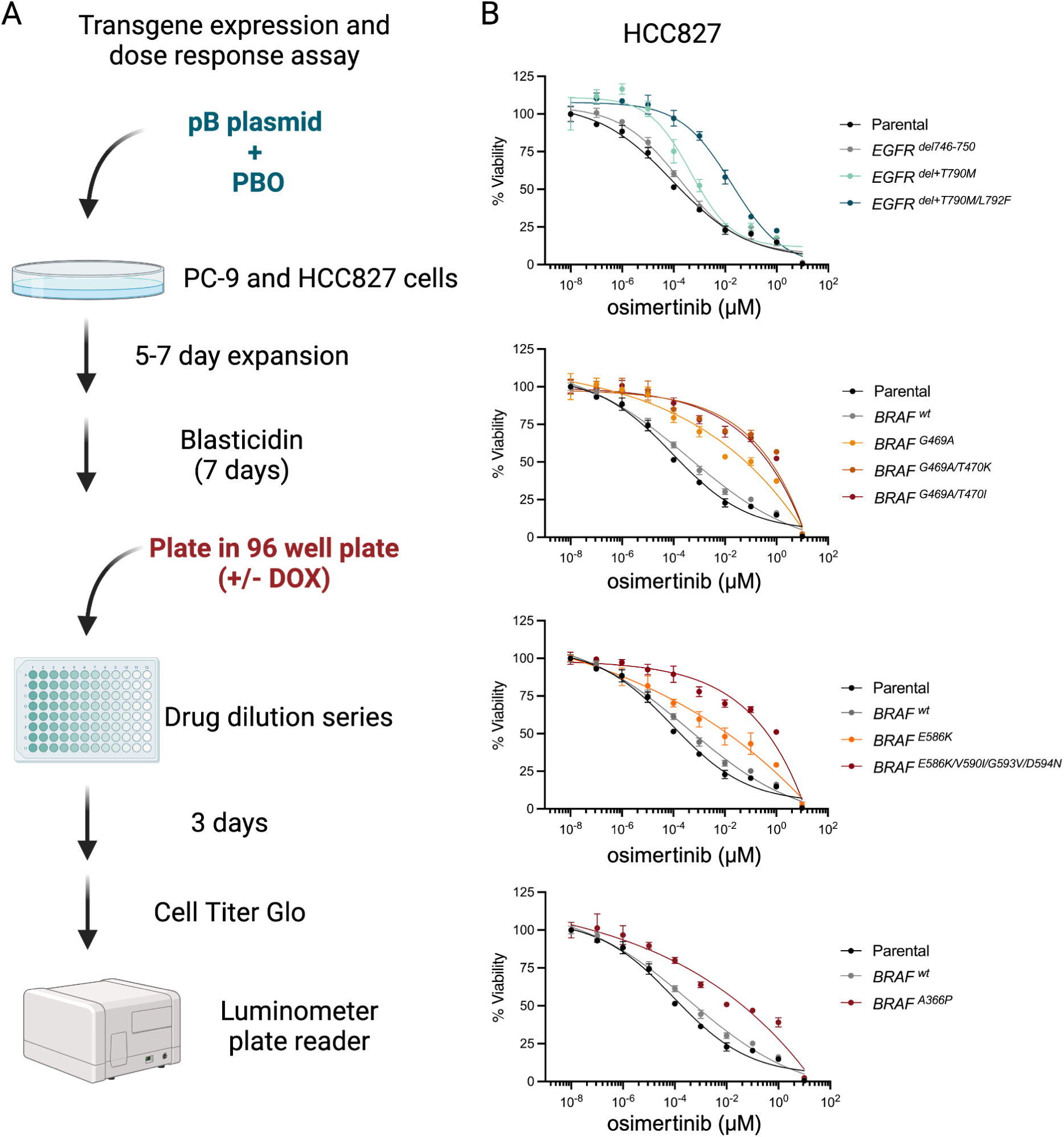
Additional information showing expression of DOX inducible *EGFR* and *BRAF* mutant transgenes recapitulates the EGFR inhibitor resistance phenotype. **A.** Schematic diagram of experimental method to generate cell lines and assay viability across a drug dilution series. **B.** Dose-response curves showing the viability of HCC827 cells expressing the specified EGFR or BRAF transgenes at escalating osimertinib concentrations.

## DISCUSSION

Tumor evolution and, consequently, drug resistance represent principal barriers to the durability of precision cancer therapies. Therefore, an ability to model and predict the numerous trajectories tumor genomes might take in order for cancers to thrive under otherwise hostile conditions is critical for generating hypotheses to drive next generation therapeutic development. Moreover, protein sequence alterations in drug resistant cells can reveal a unique view into structure-function relationships within the target protein or across binding partners and associated networks. Beyond direct sequencing of patient-derived samples or resistant cells created in vitro, a number of genetic screening technologies have become available to speed the pace of resistance variant discovery. Of particular interest are those that enable in situ genetic modification, like CRISPR base editing or prime editing. After comparing several CRISPR base editing technologies, we identified both a platform and experimental parameters that created high levels of diversity across a broad target window, for multiplexed generation of drug-resistant variants in a pooled screening context.

Relative to previously-described methods, the AID-dCas9 toolset detailed in this study produces single nucleotide and compound variants with increased efficiency and diversity, resulting in loci with a broad range of amino acid changes per targeted site. Using this platform and resistance to first (erlotinib) and third (osimertinib) generation EGFR inhibitors in EGFR mutant NSCLC as our model system, we were able to rediscover several variants in *cis* (EGFR) and *trans* (BRAF) observed in clinical isolates. Despite being an incomplete assessment of the possible genes and variants involved in resistance to these drugs, our results correspond well with clinical observations. EGFR variants (particularly T790M) explain the bulk of resistance to first generation inhibitors, whereas up to 50% of resistance mechanisms to osimertinib remain unknown and detached from EGFR mutations (8, 9). Similarly, we observed predominantly EGFR-linked variants in high dose erlotinib-resistant cells and BRAF-linked variants in the high-dose osimertinib condition, potentially reflecting differences in the mechanism of action of these two drugs. Additionally, highly abundant variants can outcompete rarer and less potent variants, particularly in the presence of strong drug selection, which may also contribute to this observed difference.

Although a number of variants identified within our screens have been found in tumor isolates, a substantial fraction either had no direct links to EGFR inhibitor resistance or, in some cases, had not yet been functionally defined. The BRAF A366P variant provides an interesting case study. Previously characterized as a rare and non-functional variant in melanoma samples, this allele validated in all in situ editing and over-expression contexts for osimertinib resistance, suggesting that it is a *bona fide* potentiator of the MAPK pathway downstream of EGFR. Separately, we discovered a series of compound variants in BRAF anchored by the well-characterized E586K mutant. While BRAF E586K on its own protected against osimertinib, these effects were much more pronounced (by logFC) when linked to V590I, G593V/C, and D594N as a compound allele. This finding also highlights the power of our AID-dCas9 platform for generating complex, multi-amino acid changes to the genome with limited biases, which is advantageous for functional assignment of known alleles, but, particularly, discovery of biologically-active compound variants.

The highly variable outcomes for each sgRNA when paired with AID-dCas9 requires unique considerations compared to similar approaches with the standard Cas9 nuclease, prime editors, or precise base editors (i.e. BE4), each of which have more predictable effects. Accordingly, we performed our screens at 1000-fold coverage, which (while sufficient for selecting the most efficiently-generated and/or potent variants) may have undersampled rare variants including those less-efficiently generated with AID-dCas9 or that reduce cell growth under basal conditions. For circumstances where recovery of such rare alleles is critical, increased coverage throughout the screening process or induction of base editing in the presence of drug could be necessary. In addition, we expected that varying the selective pressure with high or low drug dose would produce a different spectrum of mutants. Indeed, in our primary screens we found that the position and diversity of enriched sgRNAs differed depending on the dose of each drug used during selection. While we focused on validation of particularly active mutants, capable of growth in relatively-high drug concentrations, for comprehensive accounting of variants that enable drug resistance, characterization of alleles across a range of drug doses would be required. Moreover, while resistance was more pronounced with the mutant alleles, we observed significantly enhanced resistance after over-expression of unmodified *EGFR* and *BRAF* ORFs in PC-9 cells; an effect that was less pronounced in the HCC827 line. This, along with our observation that the G469A variant can synergize with T470I/K in *cis* within HCC827 but not PC-9 cells, highlights both the concern for artifacts due to transgene over-expression and also the need to validate hits with complementary approaches in distinct cellular backgrounds.

Looking forward, several technical enhancements can be made to expand the potential for variant discovery and functional assignment with the AID-dCas9 platform. As an example, our study did not identify several known drug-resistant EGFR and BRAF variants in our study (*e.g.* EGFR C797S and BRAF V600E) either due to the lack of a suitable PAM site for optimal positioning of the editing window or the inability of the AID-dCas9 base editor to produce these variants (*e.g.* T>A or G>C for EGFR C797S and T>A for BRAF V600E). In this study, we relied on the well-characterized *Staphylococcus pyogenes* (Sp)Cas9, which uses an NGG PAM. The use of a less-restrictive Cas9 (or similar), for example Cas9-NG or SpG, would enable greater density of tiling for arrayed screens and access to sites that would otherwise be missed with SpCas9 (79). While AID was capable of generating a variety of nucleotide changes, the target bases are limited to Cs and Gs (on opposing strands). Combining AID with a similarly-mutagenic ABE would substantially expand the opportunity for variant production and discovery. Due to the potential for genotoxicity from these combined editors, careful measures should be taken to limit the timing of BE expression. Recently, the use of single cell long-read and amplicon sequencing has made it possible to evaluate both genomic and transcriptomic changes after base editing (70, 80). While these approaches are currently limited by depth of sequencing at each locus, it opens the door to future variant discovery screens with transcriptional readouts. As sequencing technologies advance, this may also permit combinatorial mutagenesis screens with multiple sgRNAs expressed in each cell, resulting in multiplexed edits in *cis* and *trans*. Lastly, the highly mutagenic nature of AID-dCas9 pairs extremely well with the need for large datasets to feed artificial intelligence/machine learning (AI/ML) algorithms, which could inform future intra- and inter-protein dynamic predictions.

## ACKNOWLEDGEMENTS

We would like to thank Oleg Mayba and Soren Warming for helpful suggestions and discussions throughout the process of completing this project.

